# *Nr2e3* functional domain ablation by CRISPR-Cas9D10A identifies a new isoform and generates Retinitis Pigmentosa and Enhanced S-cone Syndrome models

**DOI:** 10.1101/2020.06.13.147785

**Authors:** Izarbe Aísa-Marín, M José López-Iniesta, Santiago Milla, Jaume Lillo, Gemma Navarro, Pedro de la Villa, Gemma Marfany

## Abstract

Mutations in *NR2E3* cause retinitis pigmentosa (RP) and enhanced S-cone syndrome (ESCS) in humans. This gene produces a large isoform encoded in 8 exons and a previously unreported shorter isoform of 7 exons, whose function is unknown. We generated two mouse models by targeting exon 8 of *Nr2e3* using CRISPR/Cas9-D10A nickase. Allele Δ27 is an in-frame deletion of 27 bp that ablates the dimerization domain, whereas allele ΔE8 (full deletion of exon 8), produces only the short isoform that lacks the dimerization and repressor domains. The Δ27 mutant shows developmental alterations and a non-progressive electrophysiological dysfunction that resembles the ESCS phenotype. The ΔE8 mutant exhibits progressive retinal degeneration, as occurs in human RP patients. Interestingly, the mutant retinas show invaginations similar to fovea-like pits. Our mutants suggest a role of *Nr2e3* as a cone-patterning regulator and provide valuable models for studying mechanisms of *NR2E3*-associated retinal dystrophies and evaluating potential therapies.

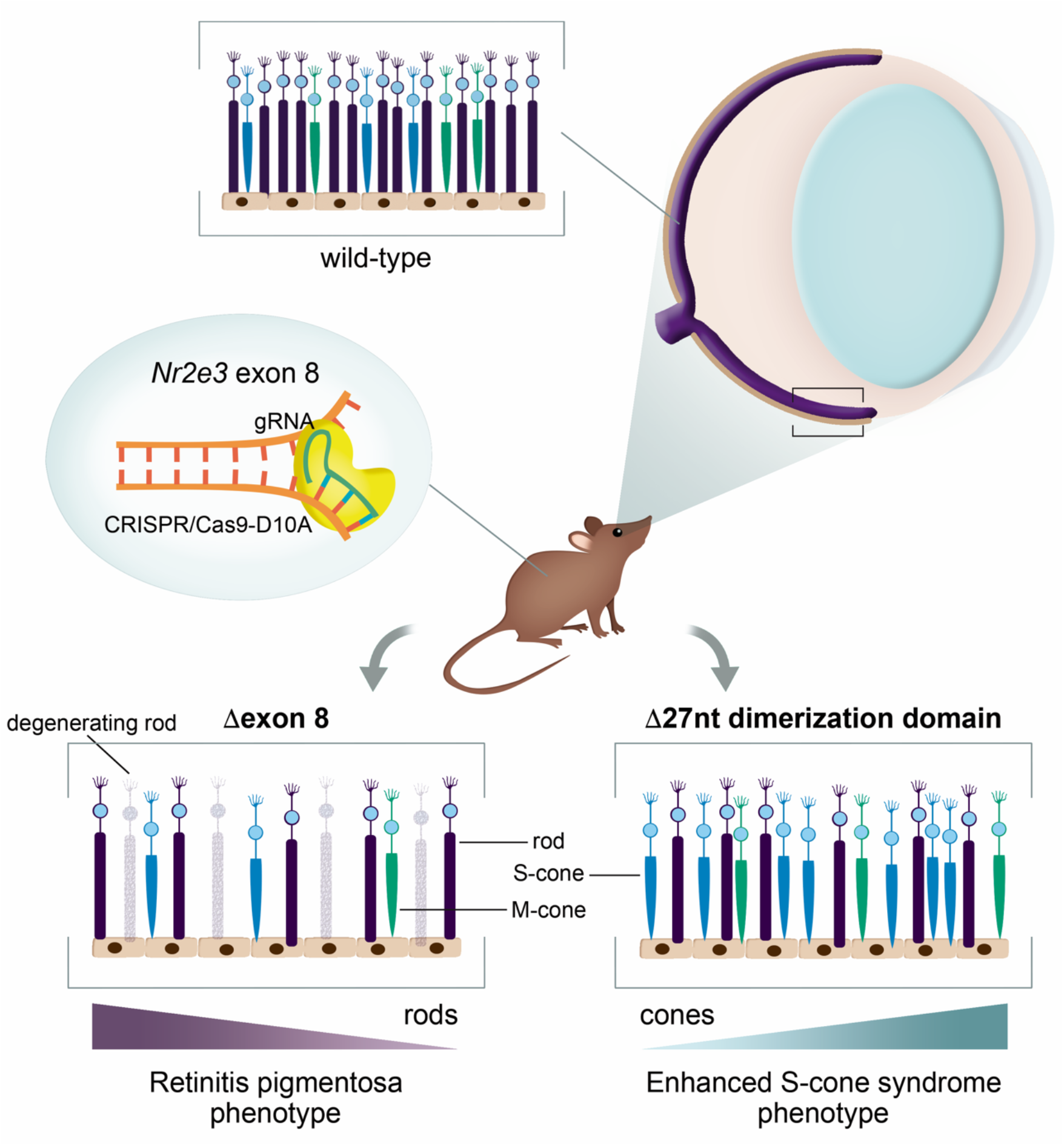

**Highlights:**

- *Nr2e3* mouse models were generated by exon 8 deletion using CRISPR/Cas9 D10A nickase.
- New *Nr2e3* mRNA retaining intron 7 encodes a short protein expressed in adult retina.
- Deletion of 9 aa of the NR2E3 dimerization domain causes enhanced S-cone syndrome.
- Deletion of exon 8 produces a phenotype similar to Retinitis Pigmentosa in mouse.

## INTRODUCTION

Inherited retinal dystrophies (IRDs) are a group of diseases associated with mutations in more than 330 genes (RetNet, the Retinal Information Network), which play critical roles in retinal function. Mutations in these genes cause alterations in retinal development or photoreceptor homeostasis, eventually leading to vision loss.

The human retina is formed by rods and three types of cones: S-cones (short wavelength), M-cones (medium wavelength) and L-cones (long wavelength), while the mouse retina is formed by rods, M-cones and S-cones. Cone photoreceptors mediate color vision and visual acuity, whereas rod photoreceptors are much more sensitive to light than cones and are excited in dim light conditions (Schnapf et al., 1987; Nathans et al., 2012).

Retinal development requires a careful orchestration of transcription factors (TFs). Retinal progenitor cells divide into post-mitotic precursor cells (PMCs), which express *Crx* to commit to the photoreceptor fate. CRX enhances the expression of photoreceptor-specific genes, among them *Nrl* and *Nr2e3*, which encode two relevant photoreceptor transcription factors (Chen et al., 1997; Freund et al., 1997). PMCs that do not express NRL follow the default pathway and differentiate into a S-cone. NRL interacts with CRX in some PMCs (Akimoto et al., 2006; Mears et al., 2001; Oh et al., 2007), thus inducing the expression of rod-specific genes and committing photoreceptor precursors to rod cell fate (Cheng et al., 2011; Peng et al., 2005).

NR2E3 (*NR2E3*; MIM# 604485), also called PNR, is an orphan nuclear receptor expressed in the retina that work synergistically with CRX and NRL to modulate the final development of rods (McIlvain and Knox, 2007; O’Brien et al., 2004; Oh et al., 2008). NR2E3 has a dual role in rod and cone fate since it can function as both a transcriptional activator of rod genes and a repressor of cone genes (Cheng et al., 2004, 2006; Haider et al., 2006). Therefore, it is necessary for cone gene inhibition and rod differentiation during retinal development (Haider et al., 2001; Cheng et al., 2004; Peng et al., 2005), and in addition, it is also relevant for photoreceptor homeostasis and is expressed at high levels in the mature retina (Haider et al., 2001; Cheng et al., 2004).

NR2E3 is a ligand-regulated transcription factor whose physiological ligands are unknown (Kobayashi et al., 1999; Qin et al., 2013). It shares the protein structure displayed by nuclear receptors consisting of a N-terminal DNA-binding domain (DBD) and a C-terminal ligand-binding domain (LBD) formed by a hydrophobic pocket of 12 helices (Mangelsdorf et al., 1995; Tan et al., 2013). The reported active form of NR2E3 is homodimeric, and dimerization is mediated by the helix 10 domain (H10) located at the LBD and close to the protein C-terminus. Structural modeling based on the LBD crystal structure revealed an auto-repressed conformation in which the AF2 region occupies the canonical cofactor binding site, which is thus required for the NR2E3 transcriptional repressor activity (Kanda and Swaroop, 2009; Tan et al., 2013). TFs are usually modulated by reversible post-translational modifications, for instance, NR2E3 sumoylation by PIAS3 is required to repress cone genes, whereas the non-sumoylated NR2E3 mainly acts as a trans-activator of rod genes (Onishi et al., 2009). In humans, this gene produces a large transcript isoform that spans 8 exons (NM_014249.4) and produces the conventional NR2E3 protein (410 aa, NP_055064.1). A shorter transcript isoform (NM_016346.4) retains intron 7 and is translated into a shorter protein of 367 aa (NP_057430.1) that lacks exon 8, which encodes the dimerization H10 and repressor AF2 domains. The physiological function of this shorter isoform is yet to be determined.

NR2E3 mutations cause either retinitis pigmentosa (RP; MIM# 268000) or enhanced S-cone syndrome (ESCS; MIM# 268100), whose most severe affectation is also named Goldmann-Favre syndrome (GFS; MIM# 26800), in both dominant and recessive forms (Coppieters et al., 2007; Escher et al., 2009; Favre, 1958; Gire et al., 2007; Haider et al., 2000; Schorderet and Escher, 2009). RP is characterized by progressive loss of rod photoreceptors, thus producing decreased peripheral vision and night vision loss. ESCS and GFS instead are characterized by an excess of S-cones in detriment of rods, most probably due to mutated NR2E3 failure in repressing cone genes during retinal development (Favre, 1958; Haider et al., 2000). Phenotype-genotype correlation is well established for the p.G65R mutation, associated with autosomal dominant RP (adRP, RP37; MIM# 611131) and characterized by the progressive degeneration of rods followed later by cones (Wright et al., 2004). However, there are no clear phenotype-genotype correlation for recessive diseases associated to *NR2E3* mutations, which suggest different disease mechanisms (Schorderet and Escher, 2009).

Animal models are extremely useful tools to study the disease mechanisms at the molecular and morphological level. The *retinal degeneration 7 (rd7*) mouse has been used as a natural model of ESCS. An insertion of a L1 retrotransposon in exon 5 of *Nr2e3* causes aberrant splicing and absence of protein in the *rd7* mouse retina (Akhmedov et al., 2000; Haider et al., 2001; Chen et al., 2006). However, some authors question *rd7* as a complete model of the human disease due to some discordances between mouse and human functional tests (Schorderet and Escher, 2009). Knockout lines of *Nr2e3* have been generated in mice (Webber et al., 2008) and zebrafish (Xie et al., 2019) resembling some of the phenotypic features found in the *rd7* retina. Even so, the molecular mechanisms causing the diverse phenotypes of *Nr2e3*-associated pathologies are still unknown.

Our group designed two mouse models to dissect *Nr2e3* function and the role of the two isoforms, and elucidate the different disease mechanisms caused by *NR2E3* mutations. To this end, CRISPR-Cas9 gene editing has been used to delete exon 8, which encodes the H10 and AF2 domains. Several modified alleles that altered the sequence of the last exon, thereby affecting the dimerization and repressor domains, were produced and we obtained homozygous mice for careful phenotypic analysis. We have thus generated two novel *Nr2e3* mouse models, which show similar phenotypic traits to human disorders, and will be very useful to study how mutations in *NR2E3* alter photoreceptor development, differentiation and survival and thus lead to either ESCS or RP diseases.

## MATERIAL AND METHODS

### Animals and ethical statement

Animal handling, euthanasia and surgical dissection was performed according to the ARVO statement for the use of animals in ophthalmic and vision research, following the guidelines for animal care of the University of Barcelona and with the approval of the Bioethics Committee (File number FUE-2019-00965313, ID 2MDLDY4WZ).

### Generation of gene-edited mice using the CRISPR/Cas9 system

The CRIPSR/Cas9 system was used to generate a *Nr2e3* mouse model by deleting the exon 8 of the locus. To minimize potential off-targets, D10A Cas9, one of the nickase mutants of Cas9, was used. Four guides were designed, two guides per deletion site, in such a way to ensure single strand breaks in the targeted acceptor site of intron 7 and 3’ UTR region of *Nr2e3* locus (Supplementary Figure 1). Several murine zygotes were injected with a number of guide-RNA and endonuclease Cas9-D10A mRNA. All embryonic procedures up to the generation of the chimaera founder mice were performed at the Mouse Mutant Core Facility, Institute for Research in Biomedicine (Barcelona, Spain). The offspring obtained was genotyped to characterize the modified alleles and to detect off-targets, if any, generated by the system. PCR products were electrophoresed in a resolutive high concentration agarose gel that allowed to resolve different size alleles without the need to perform T7 Endonuclease I Assay. When a specific band for the CRISPR-deleted allele was identified, the deletion was validated by Sanger sequencing. To confirm that the CRISPR-Cas9 technique did not introduce any off-target deletion/mutation, all the potential off-target sequences determined by a prediction software (up to three mismatches with sgRNA) were analyzed. Subsequently, the mice bearing alleles of interest were selected and crossed to obtain murine heterozygous and homozygous lines for the different mutations. Mutants with a complete or partial deletion of the exon 8 and 3’-UTR regions were selected (Supplementary Figure 2).

### Genotyping

Mouse genomic DNA was isolated from ear biopsies following overnight digestion at 55°C in a lysis buffer with proteinase K. DNA amplification by PCR was used to genotype the mouse colony. Primer pairs were used to discern between *Nr2e3* Δ27 allele (*Nr2e3* intron 7 Fw and exon 8 Rv), ΔE8 allele (*Nr2e3* intron 7 FW and down Rv), and WT alleles (Supplementary Table 1).

### RNA isolation, cDNA synthesis and Reverse Transcriptase PCR (RT-PCR)

WT mouse retinas were homogenized using a Polytron PT1200E homogenizer (Kinematica, AG, Lucerne, Switzerland). Total RNA was isolated using the RNeasy mini kit (Qiagen, Germantown, MD), following the manufacturer’s instructions with minor modifications (treatment with DNAse I during 1h). Reverse transcription reactions were carried out using the qScriptTM cDNA Synthesis Kit (Quanta BioSciences, Inc, Gaithersburg, MD). Specific primers for amplification were designed and optimized: same forward but different reverse primers were used to detect either *Nr2e3* long isoform (*Nr2e3* ex6-7 Fw and exon 8 Rv) or *Nr2e3* short isoform (*Nr2e3* ex6-7 Fw and ex7-int7 Rv) (Supplementary Table 1). RT-PCR was performed according to standard thermocycling conditions.

### Protein modeling

Protein modeling was based on the human NR2E3 structured published in the National Eye Institute (NEI) commons (https://neicommons.nei.nih.gov). Swiss-model (https://swissmodel.expasy.org) was used to align the mouse NR2E3 protein sequence to the human NR2E3 structure taking advantage of the alignment mode. The resulted mouse NR2E3 structure and the location of the different domains were visualized and analyzed using PyMOL software (The PyMOL Molecular Graphics System, Version 1.2r3pre, Schrödinger, LLC).

### Cell culture, transient transfection and expression vectors for Bioluminiscence Resonance Energy Transfer (BRET) Assays

For BRET assays, sequences encoding fusion proteins consisting of the wild type and the two mutants of *Nr2e3* fused to Renilla luciferase (Rluc) and to yellow fluorescent protein (YFP) on the C-terminal end were obtained using pRluc-N1 and pEYFP-N1 generously provided by Dr. Francisco Ciruela (Ciruela and Fernández-Dueñas, 2015). *Nr2e3* wild type, ΔE8 and Δ27 coding sequences were amplified by PCR using primers containing the restriction enzyme sites at the 5’-end of Rluc and YFP proteins, lacking the stop codon, obtaining the correct in-frame fusion proteins. Primers used can be found in Supplementary Table 1.

Human embryonic kidney (HEK293T) cells were grown in Dulbecco’s modified Eagle’s medium (DMEM) supplemented with 2 mM L-glutamine, 100 μg/ml sodium pyruvate, 100 units/milliliter penicillin/streptomycin, MEM non-essential amino acid solution (1/100), and 5% (v/v) heat-inactivated fetal bovine serum (FBS) (all supplements were from Invitrogen). Transient transfection was developed with PEI method and BRET assays were performed as previously described (Reyes-Resina *et al*., 2020). Data were fitted to a nonlinear regression equation, assuming a single-phase saturation curve with GraphPad Prism software. BRET is expressed as milliBRET units (mBU).

### Immunostaining of whole mount retinas and cone counting

Mice were sacrificed by cervical dislocation, the eye was enucleated immediately after death and placed in 1X PBS. A small hole in the cornea was performed with a needle to allow a 4% PFA solution (in 1X PBS) entering into the eye for 10 minutes. The iris was cut to remove the cornea and the lens. The whole retina was dissected from the pigmented epithelium and other ocular structures by applying gently pressure with the Dumont forceps. Careful cuts at the edges were performed to flatten the tissue and the retina was placed with the photoreceptors upside. The retina was transferred to a slide, fixed using 4% PFA in 1X PBS for an hour and washed 3 times in 1X PBS for 10 minutes each at room temperature. A solution of 0.1% Triton-X-100 and 5% sheep serum in PBS was used to block the non-specific sites and permeabilize the tissue for 1 hour. Alexa Fluor 647-conjugated PNA was incubated for 2 hours at room temperature (antibodies and dilutions are specified in Supplementary Table 2). Samples were mounted using Fluoprep (BioMerieux, Marcy-l’Étoile, France) and a coverslide, and stored at 4°C. Images were obtained by confocal microscopy (SP2, Leica Microsystems) and analyzed by ImageJ software. The number of cones per area was quantified in regions of interest (ROI) placed randomly across the whole retina. Three ROIs of the same area (6.543,895 square microns) were placed in each picture of the retina, and we took 10-15 pictures per retina. For statistical analysis R studio software was used. The Two-Way ANOVA was applied when tests for equal standard deviation (SD) and normal distribution rendered positive results.

### Immunostaining of mouse retinal sections

The procedure for dissecting the mouse neuroretina and eyecup was previously described (Toulis et al., 2016). Cryosections of mouse retinas (10-12 μm slides) were obtained using a Leica CM3050-S cryostat. Slides were thawed at room temperature for 10 minutes, washed in X1 PBS for 5 minutes and incubated in PBST (0.5% Triton-X-100 in PBS) for 15 minutes. Then, they were washed 3 times in 1X PBST for 5 minutes at room temperature and incubated in blocking buffer (5% sheep serum in 1X PBS) for 2 hours. All primary antibodies were incubated at 4°C overnight. After 3 washes in 1X PBST for 5 minutes each at room temperature, PNA, the pertinent secondary antibody and DAPI (1:300 blocking buffer, Roche) were added and incubated for 2 hours at room temperature (antibodies and dilutions are specified in Supplementary Table 2). Slides were washed 3 times in 1X PBST for 10 minutes each at room temperature and mounted using Fluoprep and a coverslide. Samples were kept at 4°C until confocal microscopy (SP2, Leica Microsystems; and Carl Zeiss LSM880). Antibodies and dilutions are specified in Supplementary Table 2.

### Transmission Electron Microscopy (TEM)

Three animals per group (12 months-old) were used. Eyes from wild type, ΔE8 and Δ27 were enucleated, the cornea was perforated using a needle to create a small hole and eyes were immersed in fixative solution (2.5% glutaraldehyde, 2% PFA in 0.1 M phosphate buffer) incubation at 4°C overnight. After several rinses (0.1 M phosphate buffer) using a shaker, eyes were post-fixed in 1% osmium tetroxide and 0,8% K_4_Fe(CN)_6_ in the dark for 2h at 4°C temperature, rinsed in double distilled water to remove the osmium. Eyes were dehydrated in ascending concentrations of acetone, and embedded in Epon (EMS). Blocs were obtained after polymerization at 60°C for 48 h. Semithin sections of 1 μm in thickness were obtained using a UC6 ultramicrotome (Leica Microsystems, Vienna, Austria), dyed with 0.5% methylene blue and observed in an optic microscope Leica DM200 (Leica Microsystems, Vienna, Austria). Ultrathin sections of 60 nm thick were obtained using a UC6 ultramicrotome (Leica Microsystems, Austria), and stained with 2% uranyless and lead citrate. Sections were observed in a Jeol EM J1010 (Jeol, Japan), and images were acquired at 80 kV with a 1k x 1k CCD Megaview camera.

### Morphometric analysis

Semi-thin sections (1 μm) for TEM (see previous subsection) of eyes from wild type, ΔE8 and Δ27 mutant mice (three animals per group, 12 months-old) were stained with 0.5% methylene blue. Sections of the central retina (containing the optic nerve) were examined and photographed under the ZOE™ Fluorescent Cell Imager (Bio-Rad, Hercules, CA, USA). The ImageJ software was used for measuring the thickness of the retinal layers at 200 μm intervals. Mann-Whitney tests were performed for statistical analysis.

### Electroretinography recordings

Young (3-4 months old) and old (7-12 months old) mice were used for each of the three genotypes (wild type, and ΔE8 and Δ27 homozygotes). Electroretinography recordings were performed in 5-8 animals per group. Dark-adapted animals were anaesthetized with an intraperitoneal injection of saline solution (NaCl 0.9%), containing ketamine (70 mg/kg) and xylazine (7 mg/kg). Before recording, pupils were dilated with one to two drops of 1% tropicamide (Alcon, Spain). To preserve the corneal surface from desiccation, a drop of 2% methylcellulose was applied (Methocel, Switzerland). Three recording electrodes (ground, reference, and corneal) were used (Burian-Allen, Hansen Ophthalmic Development Lab, Coralville, IA). In all experiments, animal handling was performed under indirect dim red light (>620 nm) and mice were kept at 37°C on a heating pad during the entire procedure. For low-intensity (−2 log Cds/m2), a single light-emitting diode was placed close to the eye. The recorded electrophysiological response was amplified and filtered (CP511 AC amplifier; Grass Instruments, Quincy, MA), and digitalized (ADInstruments Ltd, Oxfordshire, UK). Rod (b-scot), mixed (a-wave and b-wave), and oscillatory potential (OP) responses were recorded sequentially under dark background conditions, and cone (b-phot) responses were recorded following light-adaptation with background white light (50 Cd/m2). To test the effect of reducing metabolic stress by illumination, animals were light-adapted for 5 min (50 Cd/m2), and then the scotopic mixed response was recorded at different times in scotopic conditions. Rod response amplitude was measured from baseline to the peak from the scotopic recordings; the a-wave amplitude was measured from baseline to the first trough from the mixed response or to the response amplitudes at 3 and 8 msec, and the b-wave and cone response amplitudes from the first trough to the peak from the mixed (b-wave) and photopic (b-phot) responses. OP was determined as the maximum amplitude between the trough and the peak of the waves. For statistical analysis, data are presented as mean ± standard deviation (SD). The significance of the differences between genotypes was determined with t-Student tests. A p value of less than 0.05 was considered statistically significant.

## RESULTS

### *Nr2e3* mutants generated by CRISPR-Cas9 gene editing on exon 8 confirm the production of an alternative isoform that encodes a shorter NR2E3 protein isoform

To dissect the physiological function of the domains encoded in exon 8 of NR2E3 and the function of the shorter protein isoform, we designed the deletion of exon 8 of *Nr2e3* using the CRISPR-Cas9 system. In order to minimize potential off-target effects, we opted for the use of the D10A Cas9 nickase, which involves the use of two guide RNAs per site (guide design and position in Figure S1). We thus microinjected the mRNA of the Cas9 nickase plus four different guide RNAs to generate double strand breaks (DSBs) at each side of *Nr2e3* exon 8, which encodes the dimerization and repressor domains (Figure 1A), in mouse zygotes. This strategy generated several modified alleles that altered the sequence of exon 8 (around one third of the resulting chimeric embryos carried at least one modified allele). The binding and recognition of the PAM site by each of the 4 RNA guides was most probably not simultaneous, and according to our results, the upstream sequences were more prone to be cut and repaired, so that most of these gene-edited alleles were only modified at the junction of intron 7 with exon 8 (Figure S2). We generated by subsequent crosses heterozygous and homozygous strains of two selected alleles, carrying a medium and a short size deletion. The ΔE8 showed a complete deletion of exon 8 and the 3’-UTR (799 nucleotides, Figures 1A and 1B, and Figure S2). The ΔE8 allele cannot encode neither the H10 (necessary for dimerization) nor AF2 (necessary for transcriptional repression) domains. We also selected the Δ27 allele, which displays an in-frame deletion of 27 nucleotides (Figures 1A and 1C, and Figure S2) that ablates the H10 domain within an otherwise unmodified exon 8 and 3’UTR.

**Figure 1.**
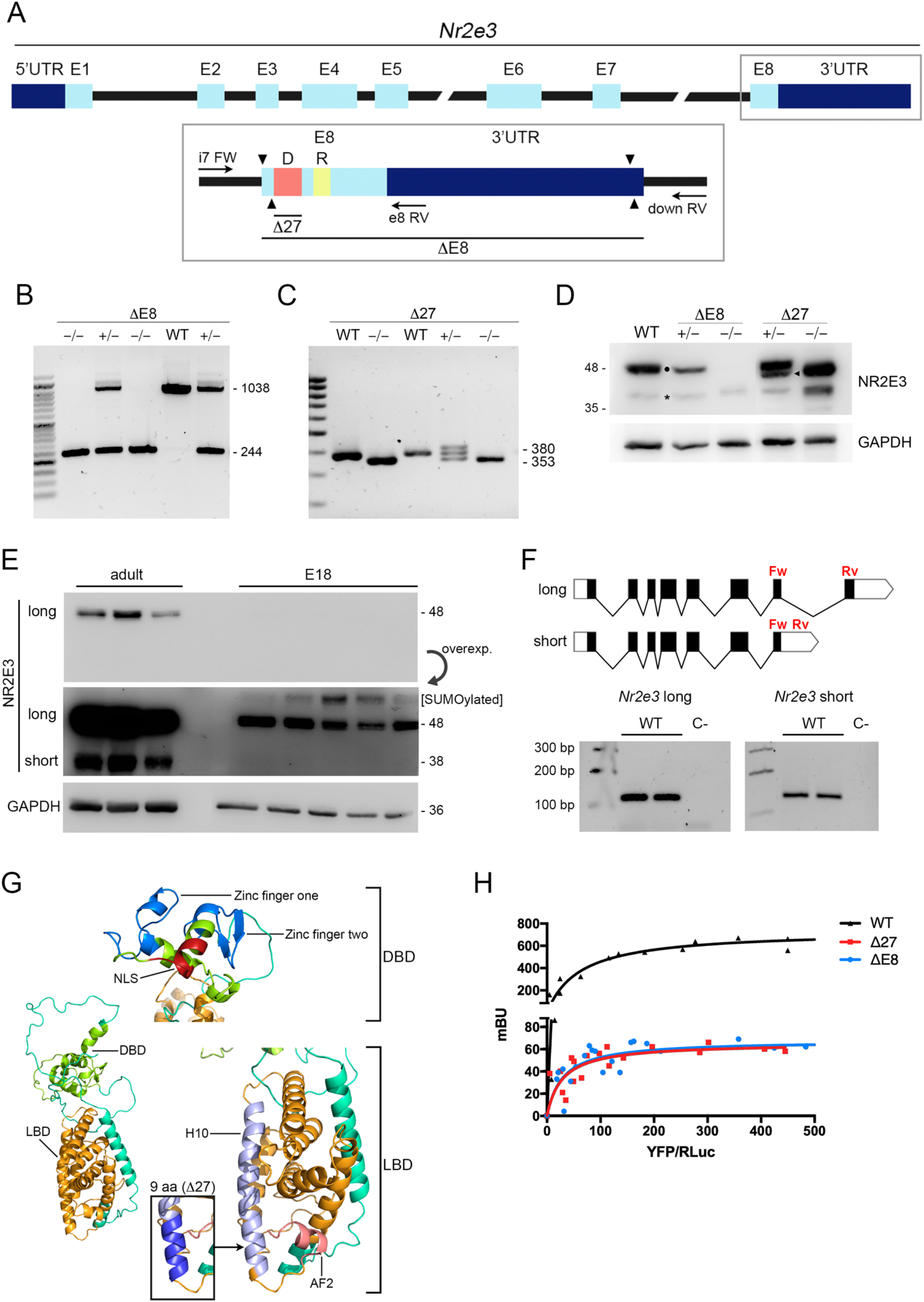
Generation of *Nr2e3* mutant alleles by gene editing with CRISPR/Cas9 D10A nickase. **A)** Schematic representation of the *Nr2e3* wild type gene with the coding exons in light blue, and 5’ and 3’UTRs in dark blue. Below, a magnification of exon 8 shows the position of the encoded dimerization (D) and repressor (R) domains. The ΔE8 mutant allele presents a full deletion of exon 8 and would result in a shorter protein without the domains encoded in exon 8. The Δ27 in-frame mutant lacks 9 amino acids that span the dimerization domain. Black arrows indicate the location of primers used for mice genotyping. Arrowheads indicate the position of the targeting sequences for the four CRISPR guide RNAs (one per each site recognized by the Cas9D10A nickases). **B)** Representative electrophoresis gel with PCR products for ΔE8 allele genotyping. The 1038 bp band corresponds to the wild type allele whereas the 244 bp band corresponds to the ΔE8 allele. **C)** PCR products for Δ27 allele genotyping. The 380 bp band corresponds to the wild type allele whereas the 353 bp band corresponds to the Δ27 allele. **D)** Immunodetection of endogenous NR2E3 isoforms in wild type and mutant mouse strains. The 48 kDa band (•) detects the NR2E3 long isoform while the 40 kDa band (*) dectects the NR2E3 short isoform. Δ27 mutant shows a slightly shorter size band of 46 kDa reflecting the in-frame deletion of 9 aa (◀). GAPDH was used as loading control. **E)** Western blot of several adult and E18 wild-type retinal protein lysates immunodetects the production of the short NR2E3 isoform in the adult retina, but not in the late embryonic stage E18 (when the mouse retina has still only cones). Note the sumoylated NR2E3 protein detected in the E18 samples, further supporting the presence of cone generepressor forms that enable the rod differentiation pathway (starting around P0). **F)** Reversetranscription PCR (RT-PCR) of adult retina mRNAs confirming the presence of transcripts for the long and short isoforms. **G)** NR2E3 protein structure showing the DNA binding domain (DBD), which contains the zinc fingers one and two and the nuclear localization signal (NLS), and the ligand binding domain (LBD), which at the C-terminus contains the H10 and AF2 domains, necessary for dimerization and repression activity, respectively. The box highlights the 9 amino acids helix deleted in the Δ27 mutant, located at the end of the H10 domain. **H)** BRET assay results for testing homodimerization of NR2E3 wild type and mutant proteins. Both Δ27 and ΔE8 mutant proteins show nearly full abrogation of NR2E3 homodimerization (one order of magnitude less than the wild-type protein).

Immunodetection by western blot of wild type mice revealed two bands for NR2E3 (Figure 1D). The band of 48 kDa corresponds to the reported full-length isoform spanning the 8 exons of *Nr2e3*. The band of 38 kDa represents a shorter isoform previously unreported in mice (only described as a transcript in humans), which retains intron 7 and produces a shorter protein isoform due to a premature STOP codon introduced early on intron 7 sequence. The ΔE8 homozygote only expresses the short isoform. The Δ27 homozygote expresses both isoforms (Figure 1D), although the longer isoform is 9 amino acids shorter. Interestingly, the short isoform is nearly 5 times more expressed in the Δ27 homozygote (~36% expression) than in the wild type animals (~8% expression). We detected a putative exonic splicing enhancer (ESE) within the 27 nucleotides deleted in the mutant, as predicted *in silico* using ESE Finder (Cartegni et al., 2003; Smith et al., 2006) (Figure S3). The predicted ESE would recruit splicing factors in the wild-type transcript, thus favoring the recognition of intron 7 and the splicing that joined exons 7 and 8. The ablation of the predicted ESE in the Δ27 homozygote would result in a favored retention of intron 7, thus increasing the expression levels of the short isoform (Figure S3).

We hypothesize that this isoform has a physiological role, since intron 7 retention has been identified in transcriptomic analysis from humans and mice. The in-frame stop codon at the very beginning of intron 7 is also evolutionary conserved in ten of the eleven vertebrate species considered (Figure S4). This isoform B displays the N-terminal transactivator domain, can bind their target motifs and potentially interact with CRX, consistent with its binding to the DBD (Peng et al., 2005; von Alpen et al., 2015), but would lack the H10 (dimerization) and AF2 (repressor) domains, which are highly conserved in all the species analyzed (Figure S4).

Remarkably, in the wild-type retina, the production of the short *Nr2e3* isoform seems restricted to the mature stage, since immunodetection in E18 embryos (when *Nr2e3* expression begins to increase (Cheng et al., 2004)) only detects the long isoform (Figure 1E). Besides, in E18 retinas, a relatively high percentage of NR2E3 is sumoylated compared to the more mature tissue, pointing to repression of cone genes (Onishi et al., 2009), in accordance to the initiation of rod differentiation (around P0) (Figure 1E). Expression of the two transcript isoforms in the adult wild-type retina was confirmed by reverse-transcription PCR using specific isoform primers (Figure 1F).

### NR2E3 proteins generated after partial in-frame or total deletion of exon 8 show impaired dimerization

Protein modelling visualizes the location of the functional domains ablated in our mutants onto the NR2E3 protein structure (Figure 1G). The DBD, containing the two zinc fingers and the nuclear localization signal (NLS), is encoded at the N-terminal part of the protein and is thus preserved in our selected alleles. Both deletions affect the LBD, which at the C-terminus displays the H10 and AF2 domains (Figure 1G).

The known functional form of full-length NR2E3 is a homodimer (Kobayashi et al., 1999), which has been observed in non-denaturing protein gel migration conditions (Roduit et al., 2009), structurally and functionally confirmed by BRET assays (Roduit et al., 2009; von Alpen et al., 2015), DNA binding assays (Escher et al., 2009; Kanda and Swaroop, 2009), as well as the crystal structure of the LBD moiety (Tan et al., 2013). Disruption of NR2E3 homodimer has been shown to impair NR2E3 repressor function (Kanda and Swaroop, 2009; Tan et al., 2013). The interaction occurs through the dimerization domain (H10 region) located in the LBD. The ΔE8 and Δ27 alleles lack, completely or partially, the H10 region. Therefore, we assayed the dimerization ability of our mutants by using BRET assay (Figure 1H), a technique that indicates a physically close molecular interaction. We fused the corresponding coding sequences of each allele and the wild-type sequence (WT) to YFP and Rluc. When expressing a constant amount of Rluc-NR2E3 WT and increasing amounts of YFP-NR2E3 WT, a saturable BRET curve was detected, indicating protein interaction. In contrast, homodimerization of the ΔE8 and Δ27 mutant proteins was greatly impaired, since BRET values were more than one order of magnitude less, thus confirming that the H10 domain is required for homodimerization of this orphan nuclear factor (Figures 1G and 1H).

### Homozygous strains of different gene-edited alleles of *Nr2e3* show severe but opposite alterations in the number of cones, and decreased number of photoreceptors

IRDs are frequently characterized by alterations in the number of photoreceptors, and most IRDs show progressive photoreceptor attrition due to a degenerative process, as it occurs in RP. In this respect, ESCS is somewhat exceptional, since it shows a gain –even if dysfunctional– in one specific type of photoreceptors, the S-cones, which are the default differentiation fate from photoreceptor precursors. Consequently, ESCS is characterized by an increase in the number of cones, even though the total number of photoreceptors is decreased, due to the reduced number of rods (Haider et al., 2000; Milam et al., 2002; Schorderet and Escher, 2009; Wright et al., 2004). Therefore, we aimed to characterize the phenotype of our mutant strains, with a special focus on cone number. We stained retinal whole mounts (>12 months-old mice, n ≥ 3) using PNA, a lectin that specifically labels cone membranes and allows the individual visualization of cones (Figure 2A). Indeed, the total cone number was statistically significantly altered when comparing the retinas of the two mutants with wild type mice (Figure 2B). The number of cones in the ΔE8 homozygous mutant retinas was 20% lower than in controls, whereas in the Δ27 was a 30% higher than in wild type retinas (Figure 2B). The Δ27 heterozygous retinas showed a slight increase (although not significant) in the number of cones, resembling an intermediate phenotype between Δ27 homozygous and wildtype retinas (data not shown). Notably, Δ27 retinas showed double cones with a thick membrane joining the twin cone cells, a trait that is frequent in some bird and fish retinas but has not been reported in mammal adult retinas (Figure 2C).

**Figure 2.**
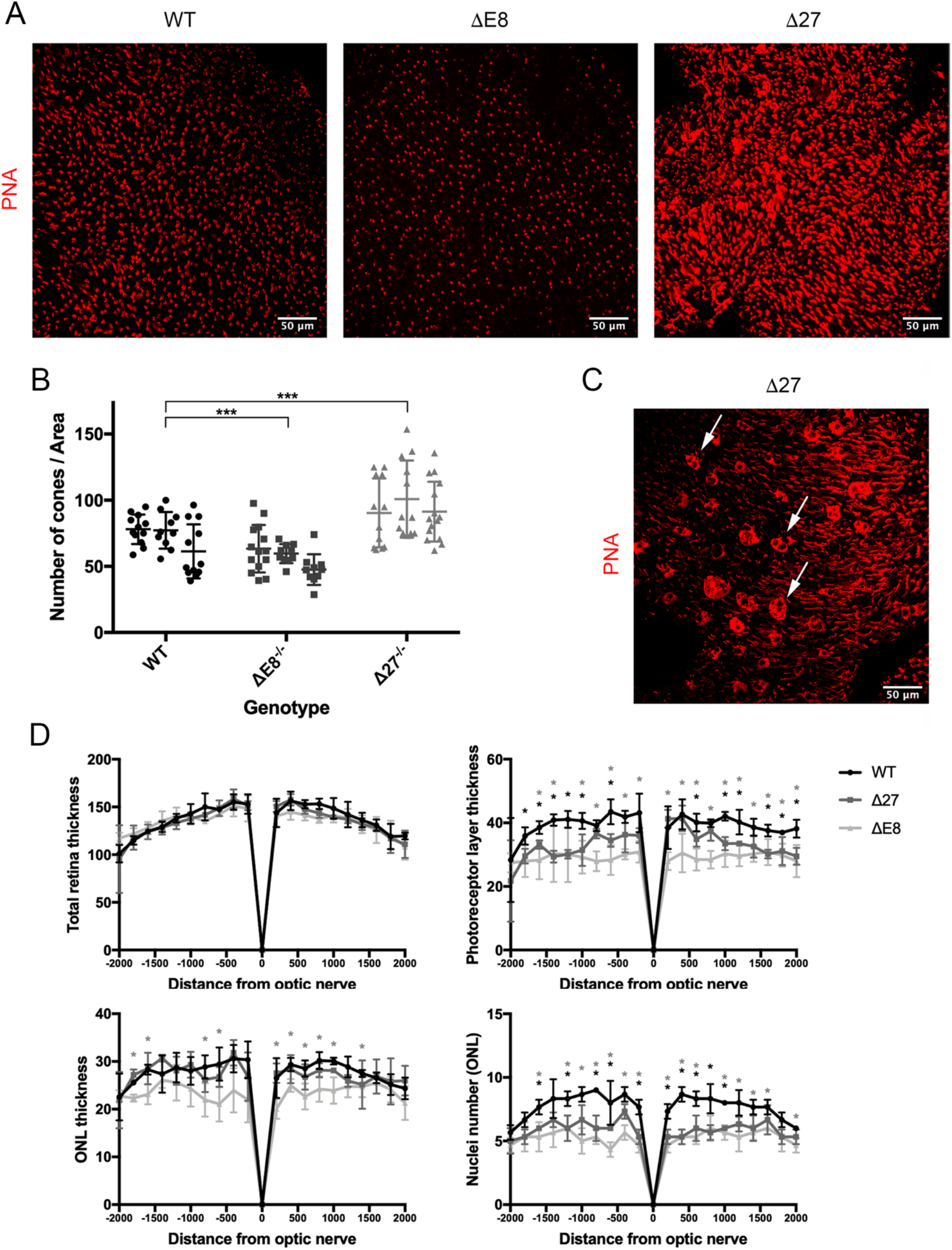
Altered number of cones and decreased number of photoreceptor nuclei in the two *Nr2e3* mutants generated by gene-editing compared to controls. **A)** Whole mount retinas from ΔE8 homozygous mice show a consistent reduction in the number of cones whereas those from Δ27 homozygous mice display a higher number of cones compared to wild-type mouse retinas. Representative images. Cones are labelled with PNA (red). **B)** Quantification of cone number showed statistically significant differences between the wildtype and homozygote retinas from the two mouse models. Mean values and standard deviation from 10–12 independent measurements in three animals per genotype were obtained and analyzed (Two-way ANOVA test, *** p<0.001). The ΔE8 mutant (low levels of truncated NR2E3 protein lacking dimerization and repressor domains) shows decreased levels of total cone photoreceptors, whereas the Δ27 mutant (normal levels of a mutant NR2E3 protein that cannot dimerize) show increased levels of total cone photoreceptors. **C)** Whole mount retinas of Δ27 homozygous mice showed structures reminiscent of twin/double cones (white arrows), frequently observed in some birds and fishes, but not in mammals. Scale bar: 50 μm. **D)** Morphometric measurements in semi-thin retinal sections containing the optic nerve show thinning of the photoreceptor layer, outer nuclear layer and number of nuclei row in mutant *Nr2e3* mice. Measurements were taken at 200 μm intervals, and mean values and standard deviation were obtained from three animals per genotype. For statistical analysis, Mann-Whitney test was performed (* p<0.05). Both ΔE8 and Δ27 mutants showed a thinner photoreceptor layer and a clear reduction of the nuclei row number of the ONL. Besides, the ONL of ΔE8 homozygous mutant mice (but not that of Δ27 mutants) was also statistically significantly thinner.

The fact that the two alleles cause an opposite effect in the number of cones suggest that the two homozygous mutant strains could mimic the phenotype of the two different retinal human diseases caused by *NR2E3* mutations. The ΔE8 phenotype might be comparable to that shown by RP, with progressive degeneration of rods and subsequent involvement of cones. The Δ27 resemble the phenotype of ESCS, with an overabundance of cones and a reduced number of functional rods.

Morphometric analysis of retinal sections, which includes morphometric parameters of retina such as cell number and thickness of different layers, has been used in examining retinal diseases.

Measurements from blue stained semi-thin sections of adult retinas at the optic nerve level were used for morphometric analysis to quantify the morphological differences observed in the ultrastructural analysis (n = 3). No clear differences in the total retina thickness were detected comparing the wild type and the mutant retinas. However, the thickness of the photoreceptor layer (PL) is decreased in both ΔE8 and Δ27 mutants, although this reduction is more evident in the ΔE8 than in the Δ27 mutant (Figure 2D upper panels). ΔE8 mutants also showed a reduction in the ONL thickness (lower number of nuclei rows), which is not observed in the Δ27 mutants. Even so, both mutants show a significative decrease in the number of nuclei present in the ONL. The decline in the number of nuclei is more notable in the ΔE8 than in the Δ27 mutant, pointing to a progressive degeneration of the retina in the ΔE8 mutant (Figure 2D lower panels).

### Photoreceptor morphology and retinal layer structure are grossly altered in the homozygous mutant retinas

To study and compare the retinal layer structure and photoreceptor morphology in our mutant models, IHC staining was performed in young (p60) and old (>12 months) wild type and mutant mice to (n = 3). Although both ΔE8 and Δ27 mutant retinas display a retinal morphology with recognizable retinal layers, the structure of the layers was obviously altered in the mutants compared to the wild type retinas (Figure 3A). However, we could not see structural differences between young and older individuals of the same genotype despite the different age. We could detect rhodopsin and cone opsin expression in the ΔE8 and the Δ27 mutant retinas. The number of cones appears particularly increased in the retinas of the Δ27 mice (Figure 3A). Remarkably, regions with bouquet-like invaginations containing the three types of photoreceptors (rods and the two type of cones) were detected in the outer retina. The outer nuclear layer (ONL) was also altered in the two mutant retinas, with a lower number of nuclei rows in the invagination fold (Figure 3A left panels). These invaginations remind of the *rd7* rosettes and whorls but are much shallower, and are detected mostly in the central retina (Figure S5). Concerning cone distribution, S-cones and M-cones are expected in opposite gradients across the wild type retina (Swaroop et al., 2010). Indeed, in wild type mouse, S-cones were present only in half of the retina. In contrast, in the Δ27 mutant, whose number of cones was increased, S-cone expression was clearly detected throughout the whole retina without a gradient distribution (Figure S5).

**Figure 3.**
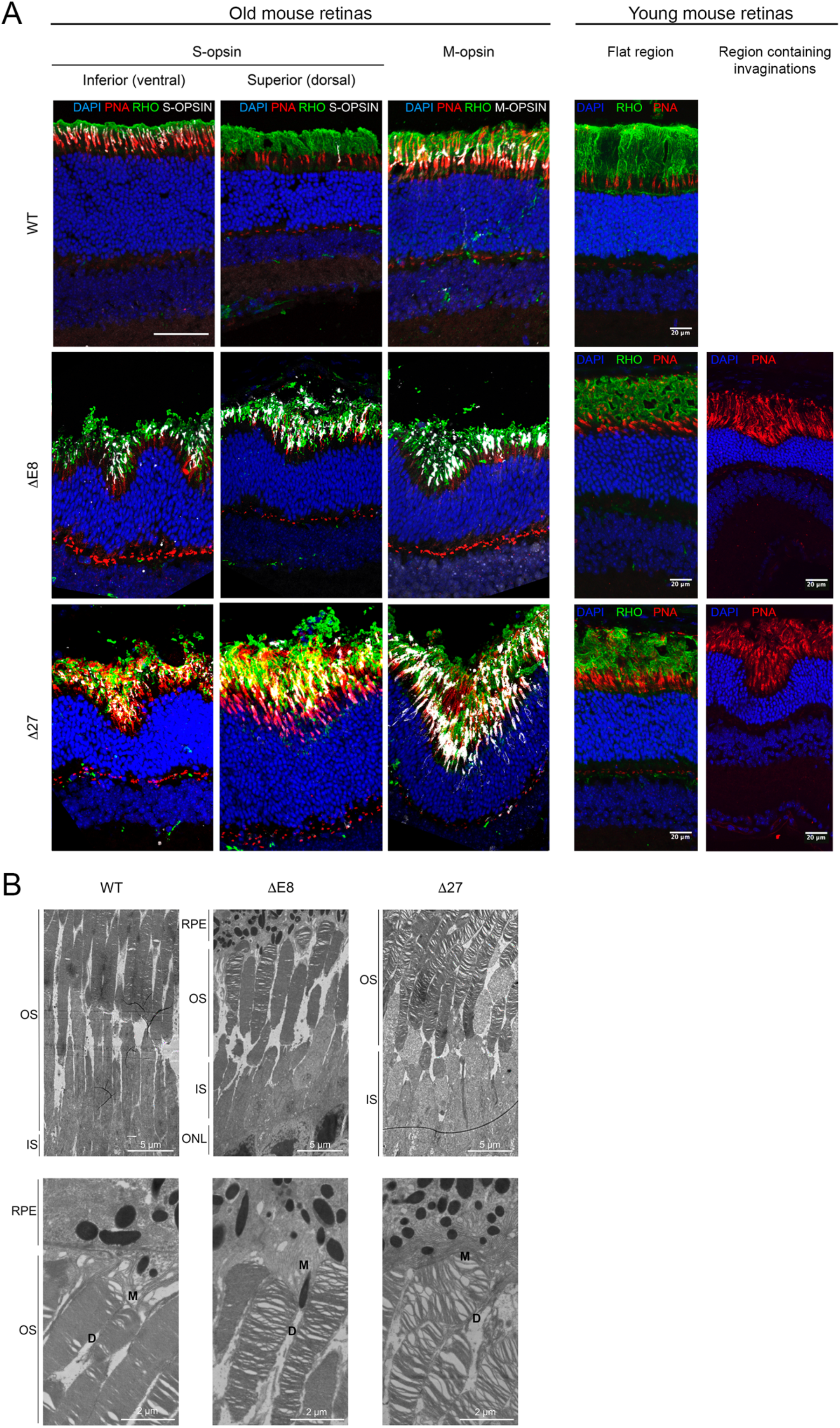
Phenotypic alterations in the retinas of the two *Nr2e3* mouse mutants include cone-rich bouquet-like invaginations and photoreceptor disk abnormalities. **A)** Immunohistochemical characterization of wild type, ΔE8 and Δ27 mutant retinas in old (>12 months) and young (p60) mice. Retinal sections were stained to all types of cones (PNA, red), rods (anti-Rho, green), and cones expressing S-or M-opsins (white). Invaginations containing rods as well as M- and S-cones were detected in Δ27 and ΔE8 homozygote retinas. The larger number of cones is particularly evident in the Δ27 retinas. Nuclei are counter stained with DAPI (blue). Scale bar: 50 μm. **B)** Ultrastructural images obtained by transmission electron microscopy (TEM) of wild type and mutant retinas, focusing on photoreceptors. A magnification of the outer segment of photoreceptors is also shown below. Retinal layers are indicated as follows: retinal pigment epithelium (RPE), outer segment (OS), inner segment (IS) and outer nuclear layer (ONL). The ΔE8 mutant shows a clear decrease in the number of photoreceptors. Both mutants display disarrayed outer segments with a clear disorganization of the photoreceptor membranous disk stacks. Scale bar: 5 μm (top) and 2 μm (bottom).

The ultrastructural analysis of photoreceptors in mutant retinas also reveals alterations in the retinal layer structure and photoreceptor morphology. The ΔE8 mutant shows a decrease in the total number of photoreceptors, in accordance with our previous results (Figure 3B upper panels).

The photoreceptor layer (PL), spanning inner and outer segments (IS and OS, respectively), is thinner compared to the wildtype and Δ27 individuals (Figure 3B upper panels). On the other hand, the Δ27 mutant presents affectation of the RPE microvilli, which apparently are more numerous than in the wild type (Figure 3B bottom panels). The ΔE8 as well as the Δ27 mutants show structural problems in disk stacking in the photoreceptor outer segments, as the membranous disks are loosely packed compared to the wild type retinas (Figure 3B, bottom panels).

### Increase of GFAP and Iba1 levels are indicative of gliosis as a response to retinal stress in the homozygous mutant retinas

These results prompted us to analyze whether the retinal alteration at the plexiform layers in the mutants was associated to retinal stress. The retina undergoes a process of degeneration of photoreceptors in response to stress and glial cells play an important role in tissue maintenance. In retinal gliosis, Müller cells (retinal macroglia cells) and microglia cells are activated and migrate upon conditions of high stress. *Nr2e3* wild type and mutant retinas were stained with GFAP and Iba1 to detect Müller cells and microglia, respectively. In the marginal regions of the retina, ΔE8 and Δ27 mutants show an increase in GFAP expression as well as more end-feet interdigitation towards the photoreceptor layer (Figure 4A top). Mutants also show Iba1-positive cells in the ONL, with an increase of microglial processes compared to the wild type, indicating stress and retinal degeneration (Figure 4A bottom).

**Figure 4.**
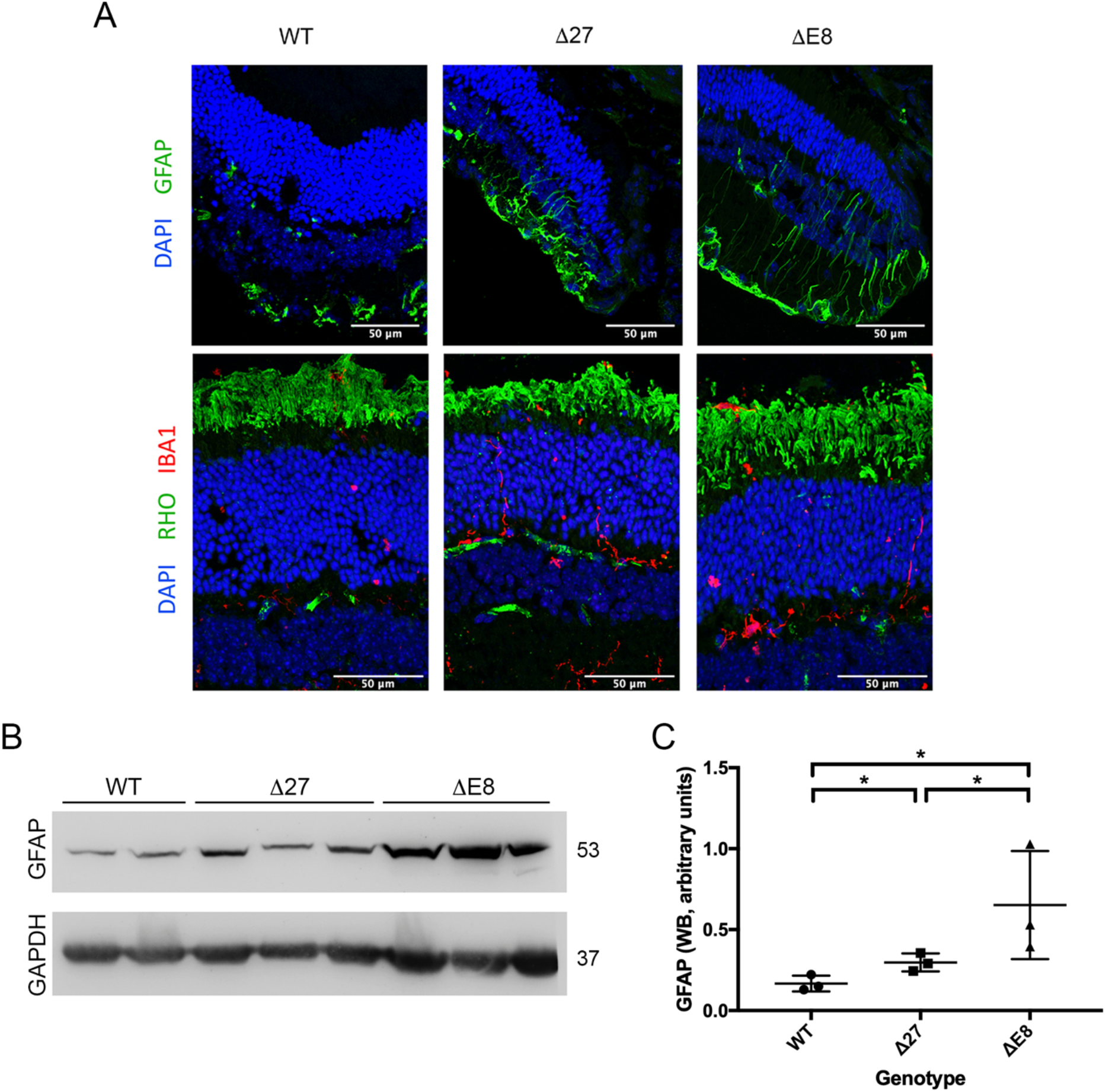
The two mutant *Nr2e3* models show an increased expression of glial markers indicative of retinal stress. **A)** Immunohistochemical staining of Müller cells (macroglia) and microglia cells in the retina. Top retinal sections were stained with anti-GFAP (mainly expressed in Müller cells, green), showing an increase in GFAP-positive processes in the two mutant *Nr2e3* models. Bottom panels show retinal sections stained for rods (Rho, green) and microglial marker (Iba1, red), showing microglia infiltration in synaptic plexiform layers and inner photoreceptor segment layer. Nuclei are counter stained with DAPI (blue). **B)** Immunodetection of GFAP (53 KDa) in the wild type and mutant retinas, and **(C)** quantification, which confirmed the statistically significant increase in GFAP-positive cells in both ΔE8 and Δ27 mutant retinas, although the levels of GFAP are higher in the ΔE8 mutant. GAPDH (37 KDa) was used as a loading control. Bars represent data (Mean ± SD) of GFAP expression (Mann-Whitney test, * p<0.05).

To compare the magnitude of the damage between the wild type and mutant individuals (n ≥ 3), GFAP protein expression was quantified by western blotting (Figure 4B). Both mutants exhibit a significative increase in the GFAP expression compared to the wild type, as expected according to the immunostaining results. However, the ΔE8 mutant shows higher levels of GFAP compared to the Δ27 mutant, indicating a more aggressive degenerative process, in accordance also with the lower number of photoreceptor nuclei observed (Figure 4C).

To assess whether reduction in the PL and ONL was due to increased cell death, we performed immunofluorescent detection of active caspase-3, an apoptosis marker. No differences were observed between the wild type and the mutant mice (data not shown).

### Retinal function is grossly altered in *Nr2e3* mutant mice

Visual function was assessed by electroretinography (ERG) waveforms obtained in 3-4 months-old and 7-12 months-old, wild type and each of the CRISPR-generated *Nr2e3* mutants (Figure 5). The electrical response to light was recorded after overnight dark-adaptation to increasing light flashes in scotopic and photopic (b-scotopic, a-mixed, b-mixed and b-photopic) conditions. The amplitude of the scotopic b-wave (b-scotopic, b-scot) assess the functionality of the rod-driven circuitry. The a-wave amplitude in these conditions (a-mixed, a-mix) reflects the activity of both rod and cone photoreceptors, whereas the b-wave (b-mixed, b-mix) include synaptic cells. Photopic b-wave (b-photopic, b-photo) test the cone-driven circuitry under photopic conditions by measuring the response to intense flashes of light in the presence of rod-saturating light stimulation.

**Figure 5.**
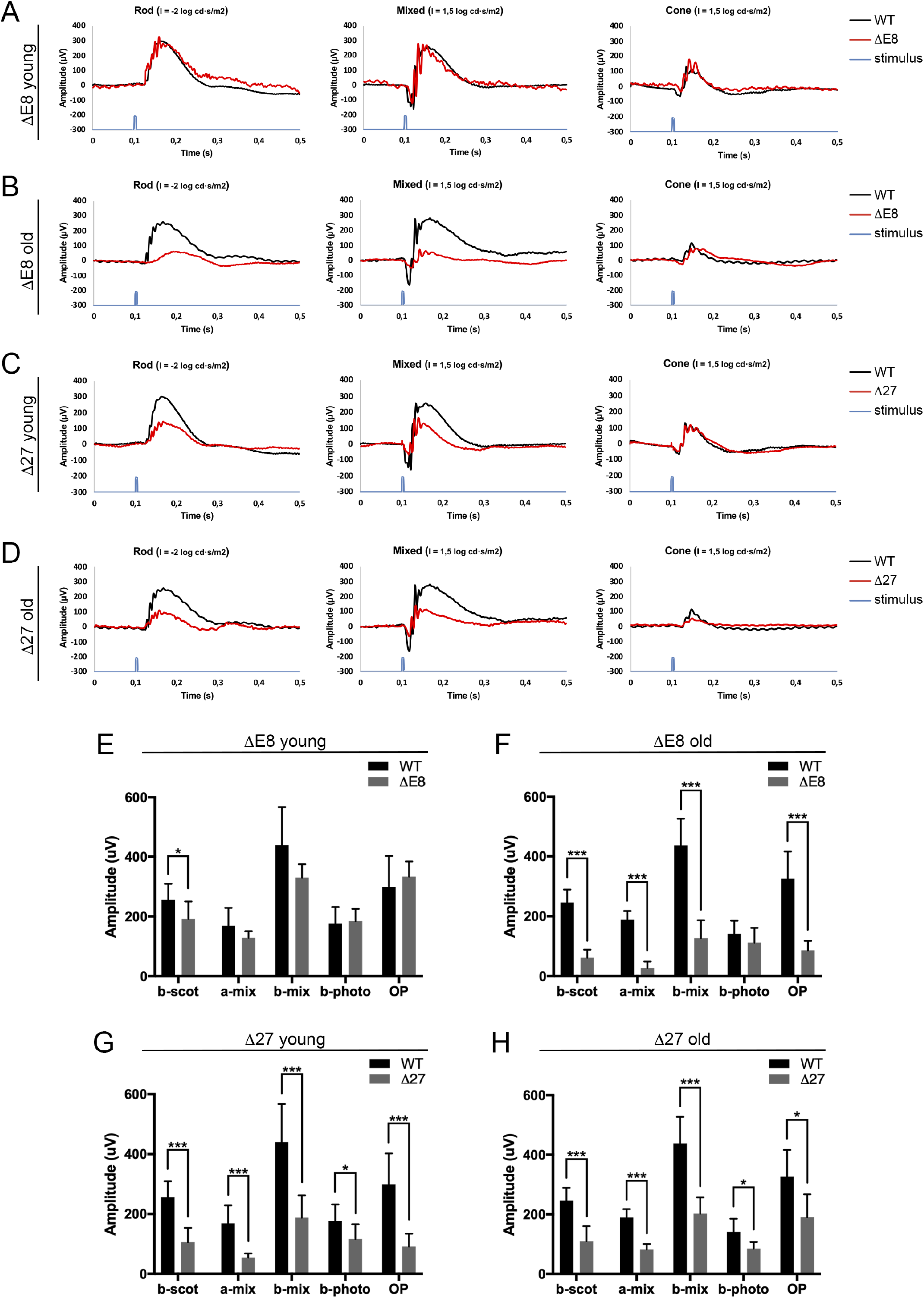
Electrophysiological recordings (ERGs) of *Nr2e3* ΔE8 and Δ27 homozygous animals show differential functional alterations in the two mutants. Representative ERG recordings obtained from wild type (black line) and *Nr2e3* mutant (red line) retinas. The two *Nr2e3* mutants show distinct recordings indicating physiological differences in the affectation of rods and cones as well as in the progression with age between the two mouse mutants compared to the control, indicating that they are potential models for different human diseases caused by *NR2E3* mutations. **A)** ERG measurements in young *Nr2e3* ΔE8 mice (3-4 months) show an initial decrease in rods activity (b-scot) compared to age-matched controls (n = 5-8 animals per group). **B)** Photoreceptor activity is clearly abrogated with age (7-12 months old), indicating a degenerative process. **C)** ERG measurements in young *Nr2e3* Δ27 mice show already alteration in photoreceptor activity (a-mix), mostly in rods (b-scot) but not so much in cones (b-photo). **D)** Remarkably, these electrophysiological alterations are stable and do not increase with age, indicating defects in the development / differentiation of photoreceptors. **E-H)** Histogram representation of the averaged ERG wave amplitude for the four animal groups: **(E)** ΔE8 young, **(F)** ΔE8 old, **(G)** Δ27 young and **(H)** Δ27 old. Amplitude of stimulus application is shown (for light stimuli details see the Material and Methods section). Bars represent data (Mean ± SD) of b-scot, a-mix, b-mix and b-photo from *Nr2e3^WTWT^* (black bars) and *Nr2e3* mutant (grey bars) mice, as indicated. Statistically significant differences (one tailed T-student test) are indicated above the histogram bars (* p<0.05, ** p<0.01, *** p<0.001).

At 3-4 months (p90-120), the amplitude of the scotopic b-wave (b-scot) showed significant differences between wild type and ΔE8 mice, indicating alterations in the rod response. No significant differences were found in the a-mix, b-mix and b-photo (Figure 5A and 5E). In contrast, at 7-12 months (p210-360), significant differences between wild type and ΔE8 mice were found when analyzing nearly all photoreceptors responses to light: in the b-scot, a-mix and b-mix recordings. However, the b-photo wave was not altered between the wild type and the ΔE8 individuals (Figure 5B and 5F). The increase in the affectation of the b-scot in young compared to old mice and the posterior addition of significant differences in the photoreceptor and synaptic cells is highly indicative of a degenerative process. The initial affectation of rods followed by alterations in other cell types is comparable to the RP human phenotype.

Remarkably, Δ27 mice showed significative differences in all of the parameters measured, b-scot, a-mix, b-mix and b-photo, at early adulthood (3-4 months old, figure 5C and 5G). These alterations were maintained and were equivalent at initial and later stages (7-12 months old, Figure 5D and 5H), thus indicating that the alterations were most probably produced by developmental defects and resembling the ESCS human phenotype.

## DISCUSSION

In humans, mutations in *NR2E3* result in functional changes in the encoded transcription factor that affect promoter target site binding, interaction with other partners, homodimerization, and regulation of transactivation/repression activity (Kanda and Swaroop, 2009; von Alpen et al., 2015), thereby altering the expression of downstream genes relevant for photoreceptor differentiation and leading to distinct retinopathies depending on the primarily affected type of photoreceptors. RP occurs in the differentiated retina as a consequence of the progressive neurodegeneration of rod photoreceptors, whereas ESCS is caused by to S-cone hypertrophy at expense of rod defective differentiation during retinal development (Coppieters et al., 2007; Escher et al., 2009; Favre, 1958; Gire et al., 2007; Haider et al., 2000; Schorderet and Escher, 2009).

We have generated different gene-edited alleles generated by CRISPR paired with the Cas9 D10A nickase. We opted for the nickase variant since we surmised that the use of two guides to produce a DSB in a precise target site plus the requirement of four guides to achieve a deletion of a medium size fragment diminished the probability of off-target events (Figures S1 and S2). At least in our hands, the Cas9 nickase was effective in generating many different alleles, although with a clear preference for one of the main target sites (34% of the mice presented gene-edited alleles at the 5’ position, with only 1,6% showing the designed deletion of the full exon 8) (Figure S2).

In this study, we present a new *Nr2e3* isoform, previously unreported in mice, generated by intron 7 retention. The fact that this transcript isoform is present at least in human and mouse, the evolutionary conservation of the in-frame stop codon at the beginning of the retained intron 7 (Figure S4), and the detection of the shorter size protein encoded in this transcript (Figure 1D), is extremely suggestive of a physiological role for this isoform. This shorter NR2E3 protein lacks both the dimerization or the transcriptional repressor domains, and it is detected in the mature but not in the late embryonic E18 retinas (Figure 1E). We believe that some of the phenotypic traits observed in our *Nr2e3* mutant mouse models might be caused by the production and/or overexpression of this isoform.

Our results suggest that the ΔE8 mutation, with the deletion of exon 8 including the 3’UTR, produces only the shorter size protein isoform, which in the adult retina is expressed at low levels. The phenotype of this mutant showed severe alterations in the maturation and homeostasis of rod photoreceptors, with progressive rod cell death, similar to the human RP phenotype. Since the expression of the short isoform does not reach the levels of the longer isoform, we cannot discard that the observed phenotype is due to both, a NR2E3 protein without the dimerization and repressor domains and the low levels of protein, which would be similar to the effect of a strong knockdown or loss-of-function allele. On the other hand, the fact that previous mouse models that ablate all isoforms of *Nr2e3* (Akhmedov et al., 2000; Haider et al., 2001; Chen et al., 2006; Webber et al., 2008) develop an ESCS syndrome instead of an RP phenotype suggests some function for the short isoform. The ΔE8 mutant presents a lower number of nuclei, and thinner PL and ONL than the Δ27 mutant (Figure 2D), indicating a more severe degeneration. The differences in the degenerative process are also significant when analyzing the markers of retinal stress, GFAP and Iba1 (Figure 4). However, we did not observed differences in the caspase-3 marker between the wild type and mutant retinas indicating degeneration is not caused by apoptosis (data not shown), in accordance with studies in the *rd7* mouse model and the knockout in zebrafish, where TUNEL assays were not significative (Cheng et al., 2011; Xie et al., 2019). Aberrant photoreceptor structure and disk packaging (Figure 3B) are very likely to alter the phototransduction pathway and affect the visual function, as shown in ERG records (Figure 5). The initial reduction in rods (b-scot) and posterior affectation of photoreceptor and synaptic activity in the ΔE8 mice is consistent with the progressive retinal neurodegeneration phenotype and attrition of photoreceptor cells observed in retinal slides. The reduction in the b-scot is indicative of higher rod affectation, similarly to what occurs in RP human patients.

In contrast, the Δ27 mutation clearly produces two proteins, the long isoform lacking the dimerization domain, and the short isoform at much higher levels than in the WT retina. The phenotypic effects observed are compatible with the inability to repress cone genes, increasing the number of S-cones and causing a very similar phenotype to ESCS in humans. The Δ27 homozygotes show electrophysiological alterations in photoreceptor and synaptic activity indicating development defects, consistent with the observed higher number of cones and similar to what occurs in ESCS patients. In humans, ESCS-associated mutations in exon 8 of *NR2E3* are also associated to a milder retinal degeneration (Audo et al., 2008). Note that although rhodopsin was detected in our mutant retinal sections, it may not indicate the presence of fully differentiated rods. Some authors have reported a rod/cone intermediate population, such “cods” have been recently characterized in retinal organoids from a NRL null patient showing an ESCS phenotype (Kallman et al., 2020). The observed phenotype of the Δ27 mutant could be the result of the short isoform overexpression plus the incapability of the long isoform to dimerize. Reported mutations (mainly missense) in *NR2E3* causing ESCS follow an autosomal recessive inheritance. The c.1101-1G>A human mutation (all reported human mutations in exon 8 are in Table S3) is a splice acceptor mutation in intron 7, which has been reported to cause ESCS in homozygosis. If intron 7 is retained, we would expect a higher expression of the short isoform, as occurs in the Δ27 mutant. Further functional tests are necessary to evaluate the pathogenic potential of the short isoform mis-regulation, which could shed light on the mechanisms underlying *Nr2e3* mutations heterogeneity.

Homodimerization is impaired in both ΔE8 and Δ27 mutants to the same extent (Figure 1H), which means that the partial and complete deletion of exon 8 disrupt similarly the dimer formation. Therefore, the 9 amino acids deleted in the Δ27 mutant are essential for homodimer formation and confirm the involvement of the alpha-helix H10 as the main dimerization domain. Although most nuclear orphan receptors act as homodimers, some function might be exerted by NR2E3 monomers binding to its recognized motifs, thus explaining why the phenotype of both mutants is different. In fact, some authors had previously reported that the dimerization potential of NR2E3 did not necessarily correlate with transcriptional activity of NR2E3 on rhodopsin and M-opsin promoters (von Alpen et al., 2015). In this context, comparison with the location and effect of patient mutations in the human ortholog gene can also shed light on the physiological role of NR2E3. The p.R385P and p.M407K mutations, respectively located in the H10 and AF2 domains (Table 1 and Table S3), result in a dimerization incompetent protein but still able to transactivate and trans-repress within a normal range (von Alpen et al., 2015). These findings strongly suggest that dimerization is not essential for the NR2E3 role as activator and/or repressor since the transcription factor would still retain the ability to bind DNA as a monomer. This hypothesis is supported by the fact that mutations located outside the dimer interface also cause typical ECSC, indicating that other factors, such as binding and recruitment of co-repressors, might be also altered mechanisms in disease.

**Table 1.**
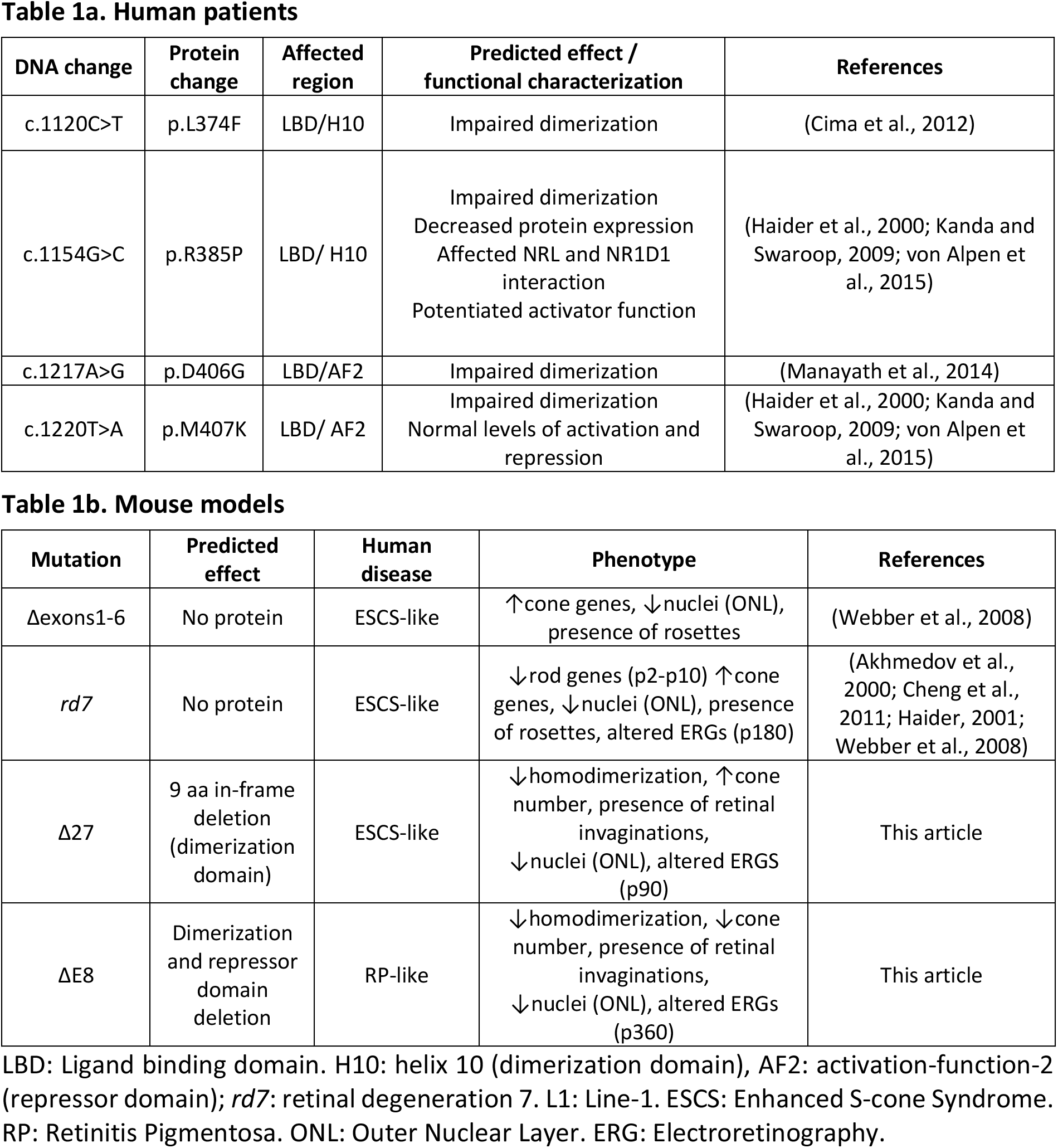
Functional effect of NR2E3 mutations in exon 8 in human and mouse.

Notably, whole mount retinas of Δ27 mice show structures compatible with double or twin cones (Figure 2C). Double cones are formed by a large cone termed the chief cone and a small one termed the accessory cone. Twin cones instead refer to two identical cones that adhere together. Double cones have been identified in some birds and fish retina, whereas twin cones have been described in fish retina. Neither double cones nor twin cones have ever been described in the mammal retina and its appearance suggest a possible role of *Nr2e3* as a regulator in cone patterning. On the other hand, the ONL and PL in the two mutants present cone-rich invaginations (Figure 3A) which are somewhat similar to the rosettes detected in the *rd7* mouse retina, although in our models the invaginations are less pronounced and are mainly detected in the central retina. In fact, these invaginations remind of fovea-like bouquets as described for the canine cone-rich (but not cone-exclusive) region in the retina centralis (Beltran et al., 2014; Kostic and Arsenijevic, 2016). The fovea, the central part of the retina responsible for visual acuity and color vision, includes a cone-exclusive tiny region called macula, in the human eye. Fovea-like regions with a high density of cones and reduced number of rods (with thinning of the ONL and a clear decrease in the number of photoreceptor nuclei) have been identified in some mammals, such as primates and dogs, but are not found in mice, whose retina is formed mostly by rods, which account to up 97% of photoreceptors (Jeon et al., 1998). It is tempting to speculate that *Nr2e3* is relevant during the retinal development not only for determining photoreceptor fate but also for retinal patterning and photoreceptor type distribution, which might impact in the formation of the fovea and macula.

In summary, we have identified two different protein isoforms produced by the *Nr2e3* gene, and generated two mouse models by deleting different domains encoded in the last *Nr2e3* exon, leading to two models of different human retinal dystrophies, RP and ECSC (Figure 6). The *Nr2e3 ΔE8* mouse model expresses only the short isoform of *Nr2e3*, which lacks the dimerization and repressor domains. *Nr2e3 ΔE8* mice present a degenerative process in mature rods, which causes high levels of stress and inflammation and leads to the progressive affectation of the remaining retinal cells. The *Nr2e3 Δ27* mouse model expresses both isoforms, but none can dimerize. This model exhibits a developmental disfunction where cone genes cannot be repressed, which results in an increase in the number of S-cones and the consequent decrease in the number of rods, thus affecting the retinal function. The two models present fovea-like cone-rich regions which may reflect the role of *Nr2e3* not only in photoreceptor fate but also in cone distribution and retinal patterning.

**Figure 6.**
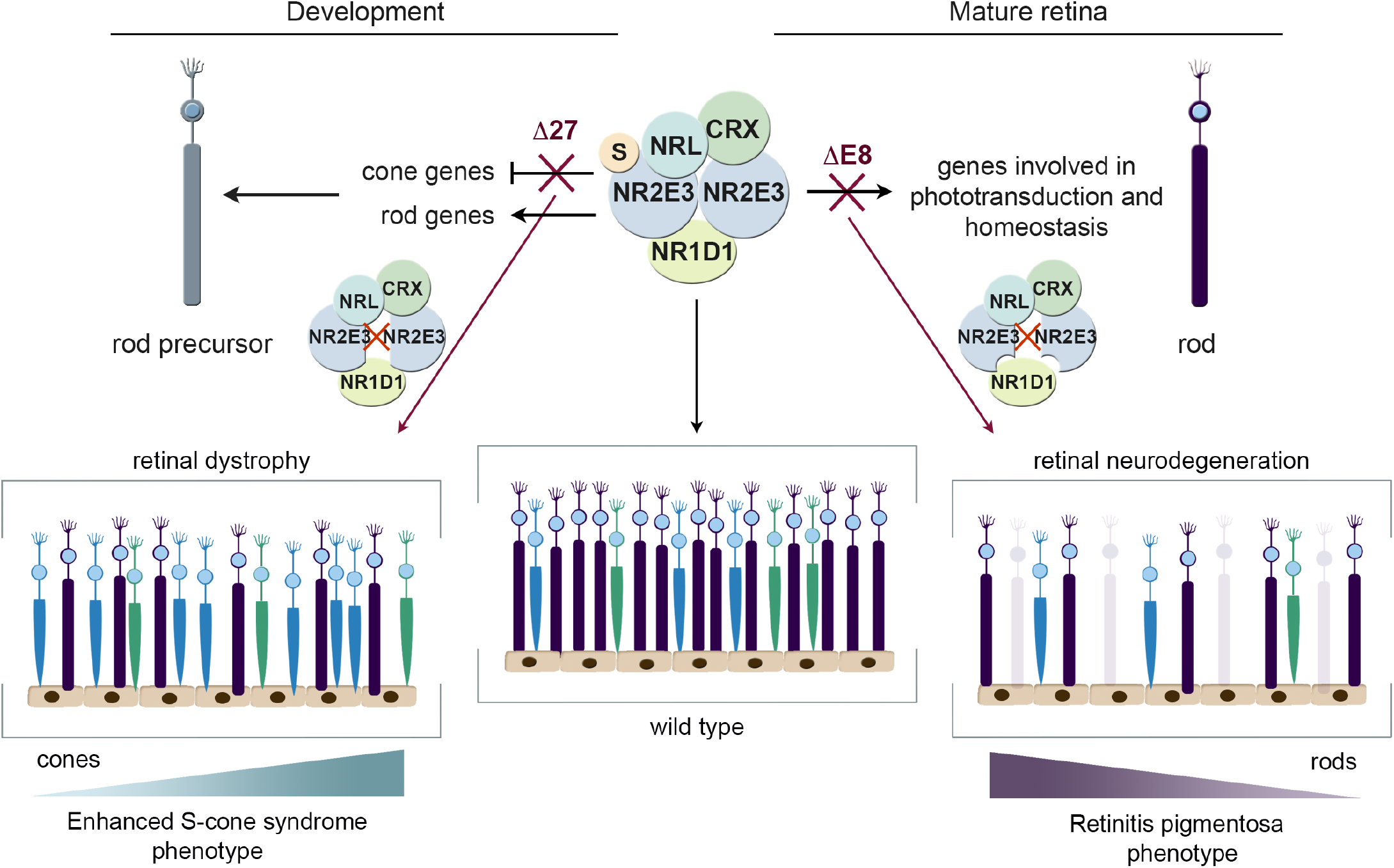
Modelling *Nr2e3*-retinal dystrophies in mouse to generate mutants resembling the enhanced S-cone syndrome (ESCS) and retinitis pigmentosa (RP) human phenotypes. NR2E3 is necessary for inhibition of cone genes (default pathway) and activation of rod differentiation genes in retinal precursors during retinal development, but it is also relevant for rod functional maintenance and survival. S indicates reversible SUMO modification of NR2E3, since sumoylation is required for cone gene repression (Onishi et al., 2009). Mutations that affect homeostasis of rod photoreceptors would cause RP, while mutations affecting repression of S-cone genes would cause ESCS. The Δ27 mutant encodes a dimerization incompetent variant of NR2E3, failing to repress cone genes leading to increased number of dysfunctional cones, triggering initial death of rods, and causing an ESCS-like phenotype, mostly reflecting a gain-of-function effect. The ΔE8 mutant lacks the dimerization and repression domains and only expresses the short isoform of *Nr2e3* at low levels, failing to maintain the homeostasis of mature rods, probably reflecting a loss-of-function effect, thus leading to a slow and progressive attrition of rods and also cones with age, and showing a phenotype similar to the RP patients.

Our *Nr2e3* ΔE8 model is the first mouse model of RP caused by mutations in *Nr2e3*. The *Nr2e3* Δ27 model concords with the knockout model in mice (Webber et al., 2008) and the *rd7* (Akhmedov et al., 2000; Haider, 2001; Chen et al., 2006), and resembles the ESCS human phenotype (Webber et al., 2008). Furthermore, the previous reported mice models for *Nr2e3* (Table 1) are knockouts, which implies that no NR2E3 protein is expressed. Our models allow us to dissect *Nr2e3* function as they lack different domains but continue to be expressed in the retina. Future research using transcriptomic and epigenomic studies will help to dissect the retinal function of the two isoforms of NR2E3 in photoreceptor differentiation and cone distribution, and how different mutations cause distinct retinal dystrophies.

Recent studies propose *Nr2e3* as a genetic modifier and broad-spectrum therapeutic agent to treat multiple forms of RP (Li et al., 2020), highlighting the importance of understanding the molecular mechanisms of *Nr2e3*-related pathways. Our models provide a valuable tool in studying *Nr2e3*-caused diseases and allow us to comprehend molecular mechanisms of disease by dissecting genetic pathways and evaluate therapeutics.

## Supporting information

Supplementary Figures with legends

## AUTHOR CONTRIBUTIONS

I.A-M. performed the experiments; M.J.L-I. generated the models and initially characterized the alleles; J.L. and G.N. performed the BRET assays; S.M and P. de V. performed the ERGs; G.M designed and supervised the experimental work, and provided the funding.

## ACKNOWLEDGEMENTS

The authors also acknowledge the technical support of Elena Laplaza and Laura Jiménez.

## FUNDING

This research was supported by grants SAF2013-49069-C2-1-R, SAF2016-80937-R (Ministerio de Economía y Competitividad/FEDER), ACCI 2015 and ACCI 2016 (CIBERER /ISCIII) and 2017 SGR 738 (Generalitat de Catalunya) to GM; La Marató TV3 (Project Marató 201417-30-31-32); and the Instituto de Salud Carlos III, cofounded with the European Regional Development Fund (ERDF) within the “Plan Estatal de Investigación Científica y Técnica y de Innovación 2017–2020” (RD16/0008/0020; FIS/PI 18-00754) to PdlV. IAM is a fellow of the APIF-2019 (Universitat de Barcelona).

## CONFLICT OF INTEREST STATEMENT

The authors declare that they have no conflict of interest.

## Abbreviations

IRDs: Inherited retinal dystrophies
PMCs: Post-mitotic cells
rd7: retinal degeneration 7
ESCS: Enhanced S-cone syndrome
RP: Retinitis Pigmentosa

## Notes

### Competing Interest Statement

The authors have declared no competing interest.

