## Supplementary Figures with legends for "*Nr2e3* functional domain ablation by CRISPR-Cas9D10A identifies a new isoform and generates Retinitis Pigmentosa and Enhanced S-cone Syndrome models"

**Supplementary Table 1. Primers used for PCR and sequencing**

| Gene | Strand | Sequence (5' – 3') | Use |
|---|---|---|---|
| mNr2e3 ATG | Fw | gaagatctATGAGCTCTACAGTGGCTGCC | Cloning using BglII restriction site |
| mNr2e3 WT | Rv | cgcaagcttGTTTTTGAACATGTCACACAG GAG | Cloning using HindIII restriction site |
| mNr2e3 delE8 | Rv | cgcaagcttCCTCACAGGCTGGCTGGG | Cloning using HindIII restriction site |
| del27 | Fw | CACGGCTGAGCGCATTGGGAACACTCCG | Site-directed mutagenesis |
| del27 antisense | Rv | CGGAGTGTTCCCAATGCGCTCAGCCGTG | Site-directed mutagenesis |
| *Nr2e3* intron 7 | Fw | AGTCCCTGTTTTCATCACGCT | Genotyping |
| *Nr2e3* exon 8 | Rv | GCTGAGAAAGGCAGGGAACTA | Genotyping |
| *Nr2e3* down | Rv | AGAGGGAAGGAAGATCCCGT | Genotyping |
| *Nr2e3* ex6-7 | Fw | CTTCAAACCTGAAACACGAGGC | RT-PCR |
| *Nr2e3* ex7-8 | Rv | TTCCCAAACCTCACAGGCTG | RT-PCR |
| *Nr2e3* ex7-int7 | Rv | GGGTCATTGCTGTCACCTCAG | RT-PCR |

Fw: Forward. Rv: Reverse

**Supplementary Table 2. Antibodies used for western blot immunodetection (WB), whole mount immunostaining (WM) and immunohistochemistry (IHC).**

| Antibody | Reference | Description* | Dilution | Blocking solution |
|---|---|---|---|---|
| α-GFP | Abcam ab290 | Rabbit pAb | 1:1000 (WB) | 10% milk |
| α-GFP | Abcam ab183734 | Mouse mAb | 1:1000 (WB) | 10% milk |
| α-alpha-tubulin | Sigma-Aldrich T5168 | Mouse mAb | 1:1000 (WB) | 10% milk |
| α-Rho | Abcam ab98887 | Mouse mAb | 1:250 (IHC) | 5% sheep serum |
| α-L/M-opsins | Millipore AB5405 | Rabbit pAb | 1:300 (IHC) | 5% sheep serum |
| α-S-opsin | Millipore AB5407 | Rabbit pAb | 1:300 (IHC) | 5% sheep serum |
| α-GFAP | Millipore MAB360 | Mouse mAb | 1:500 (WB), 1:200 (IHC) | 5% sheep serum |
| α-Iba1 | Abcam ab178846 | Rabbit mAb | 1:200 (IHC) | 5% sheep serum |
| α-NR2E3 | Abcam ab41922 | Mouse mAb | 1:1000 (WB) | 10% milk |
| α-NR2E3 | R&D Systems H7223 | Mouse mAb | 1:1000 (WB) | 10% milk |
| PNA Alexa Fluor 647 conjugate | Thermo Fisher Scientific L32460 | Peanut agglutinin | 1:50 (WM, IHC) | 5% sheep serum |
| Alexa Fluor 568 α-Rabbit IgG | Thermo Fisher Scientific A11011 | Goat pAb | 1:300 (IHC) | 5% sheep serum |
| SAMPO: α-mouse igG-PO | Sigma-Aldrich A-5906 | Sheep pAb | 1:2000 (WB) | 10% milk |
| DARPO: α-rabbit IgG-PO | GE Healthcare NA934VS | Donkey pAb | 1:2000 (WB) | 10% milk |

* mAb: Monoclonal Antibody. pAb: Polyclonal Antibody

**Supplementary Table 3. Table 1. List of NR2E3 human mutations located in exon 8, classified by molecular effect, with nucleotide and protein changes and associated clinical phenotype.**

| Mutation type | DNA change | Protein change | Zygosis | Inheritance | Reported phenotype | Ref. |
|---|---|---|---|---|---|---|
| Missense | c.1112T>G | p.L371W | HT | AR | ESCS | (Ripamonti et al., 2014) |
| Missense | c.1118T>C | p.L373P | HT | AR | ESCS | (Minnella et al., 2018; Murro et al., 2019) |
| Missense | c.1120C>T | p.L374F | HT | AR | ESCS | (Cima et al., 2012) |
| Missense | c.1127C>T | p.P376L | HT | AD/AR | Retinal dystrophy | (Li et al., 2016) |
| Missense | c.1154G>C | p.R385P | HT | AR | ESCS | (von Alpen et al., 2015; Haider et al., 2000; Kanda and Swaroop, 2009) |
| Missense | c.1217A>G | p.D406G | HM | AR | GFS | (Manayath et al., 2014) |
| Missense | c.1220T>A | p.M407K | HM | AR | ESCS | (von Alpen et al., 2015; Haider et al., 2000; Kanda and Swaroop, 2009) |
| Missense | c.1225A>G | p.K409E | HT | AR | Retinal dystrophy | (Murro et al., 2019) |
| Splicing | c.1101-1G>A | Aberrant splicing | HM | AR | ESCS | (Audo et al., 2008; Schorderet and Escher, 2009) |
| Small deletion | c.1194delT | p.P399Qfs*44 | HT | AR | ESCS | (Hull et al., 2014) |
| Small deletion | c.1223delT | p.F408Sfs*70 | HT | AR | Retinal dystrophy | (Murro et al., 2019) |

HM: Homozygosis. HT: Heterozygosis. AR: Autosomal recessive. AD: Autosomal dominant. ESCS: Enhanced S-cone syndrome. GFS: Goldmann-Favre syndrome (a severe stage of ESCS). From Human Gene Mutation Database (HGMD) http://www.hgmd.cf.ac.uk/ac/index.php

**REFERENCES Suppl. Table 3**


von Alpen, D., Tran, H.V., Guex, N., Venturini, G., Munier, F.L., Schorderet, D.F., Haider, N.B., and Escher, P. (2015). Differential dimerization of variants linked to enhanced S-cone sensitivity syndrome (ESCS) located in the NR2E3 ligand-binding domain. Hum. Mutat. *36*, 599–610.

Audo, I., Michaelides, M., Robson, A.G., Hawlina, M., Vaclavik, V., Sandbach, J.M., Neveu, M.M., Hogg, C.R., Hunt, D.M., Moore, A.T., et al. (2008). Phenotypic Variation in Enhanced S-cone Syndrome. Invest. Ophthalmol. Vis. Sci. *49*, 2082–2093.

Cima, I., Brecelj, J., Sustar, M., Coppieters, F., Leroy, B.P., De Baere, E., and Hawlina, M. (2012). Enhanced S-cone syndrome with preserved macular structure and severely depressed retinal function. Doc. Ophthalmol. *125*, 161–168.

Haider, N.B., Jacobson, S.G., Cideciyan, A. V., Swiderski, R., Streb, L.M., Searby, C., Beck, G., Hockey, R., Hanna, D.B., Gorman, S., et al. (2000). Mutation of a nuclear receptor gene, NR2E3, causes enhanced S cone syndrome, a disorder of retinal cell fate. Nat. Genet. *24*, 127–131.

Hull, S., Arno, G., Sergouniotis, P.I., Tiffin, P., Borman, A.D., Chandra, A., Robson, A.G., Holder, G.E., Webster, A.R., and Moore, A.T. (2014). Clinical and Molecular Characterization of Enhanced S-Cone Syndrome in Children. JAMA Ophthalmol. *132*, 1341–1349.

Kanda, A., and Swaroop, A. (2009). A comprehensive analysis of sequence variants and putative disease-causing mutations in photoreceptor-specific nuclear receptor NR2E3. Mol. Vis. *15*, 2174–2184.

Li, N., Ding, Y.U., Yu, T., Li, J., Shen, Y., Wang, X., Fu, Q., Shen, Y., Huang, X., and Wang, J. (2016). Causal variants screened by whole exome sequencing in a patient with maternal uniparental isodisomy of chromosome 10 and a complicated phenotype. Exp. Ther. Med. *11*, 2247–2253.

Manayath, G.J., Namburi, P., Periasamy, S., Kale, J.A., Narendran, V., and Ganesh, A. (2014). A novel mutation in the NR2E3 gene associated with Goldmann-Favre syndrome and vasoproliferative tumor of the retina. Mol. Vis. *20*, 724–731.

Minnella, A.M., Pagliei, V., Savastano, M.C., Federici, M., Bertelli, M., Maltese, P.E., Placidi, G., Corbo, G., Falsini, B., and Caporossi, A. (2018). Swept source optical coherence tomography and optical coherence tomography angiography in pediatric enhanced S-cone syndrome: a case report. J. Med. Case Rep. *12*, 287.

Murro, V., Mucciolo, D.P., Sodi, A., Passerini, I., Giorgio, D., Virgili, G., and Rizzo, S. (2019). Novel clinical findings in autosomal recessive NR2E3-related retinal dystrophy. Graefe’s Arch. Clin. Exp. Ophthalmol. *257*, 9–22.

Ripamonti, C., Aboshiha, J., Henning, G.B., Sergouniotis, P.I., Michaelides, M., Moore, A.T., Webster, A.R., and Stockman, A. (2014). Vision in Observers With Enhanced S-Cone Syndrome: An Excess of S-Cones but Connected Mainly to Conventional S-Cone Pathways. Invest. Ophthalmol. Vis. Sci. *55*, 963–976.

Schorderet, D.F., and Escher, P. (2009). NR2E3 mutations in enhanced S-cone sensitivity syndrome (ESCS), Goldmann-Favre syndrome (GFS), clumped pigmentary retinal degeneration (CPRD), and retinitis pigmentosa (RP). Hum. Mutat. *30*, 1475–1485.

A

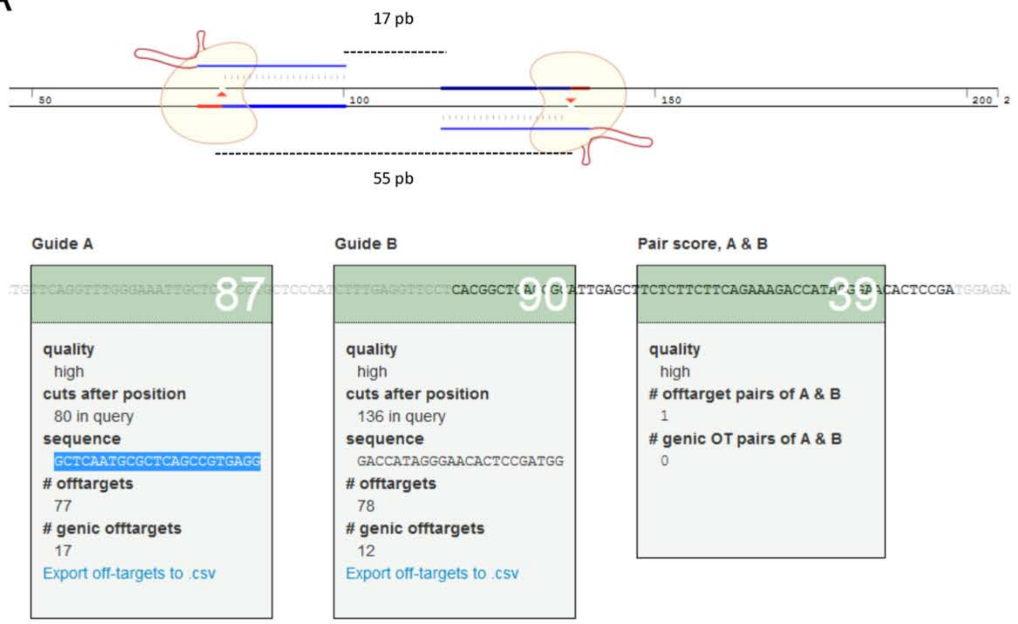


Guides 5':
gRNA NR2e3 51 GCTCAATGCGCTCAGCCGTG
gRNA NR2e3 52 GACCATAGGGAACACTCCGA

B

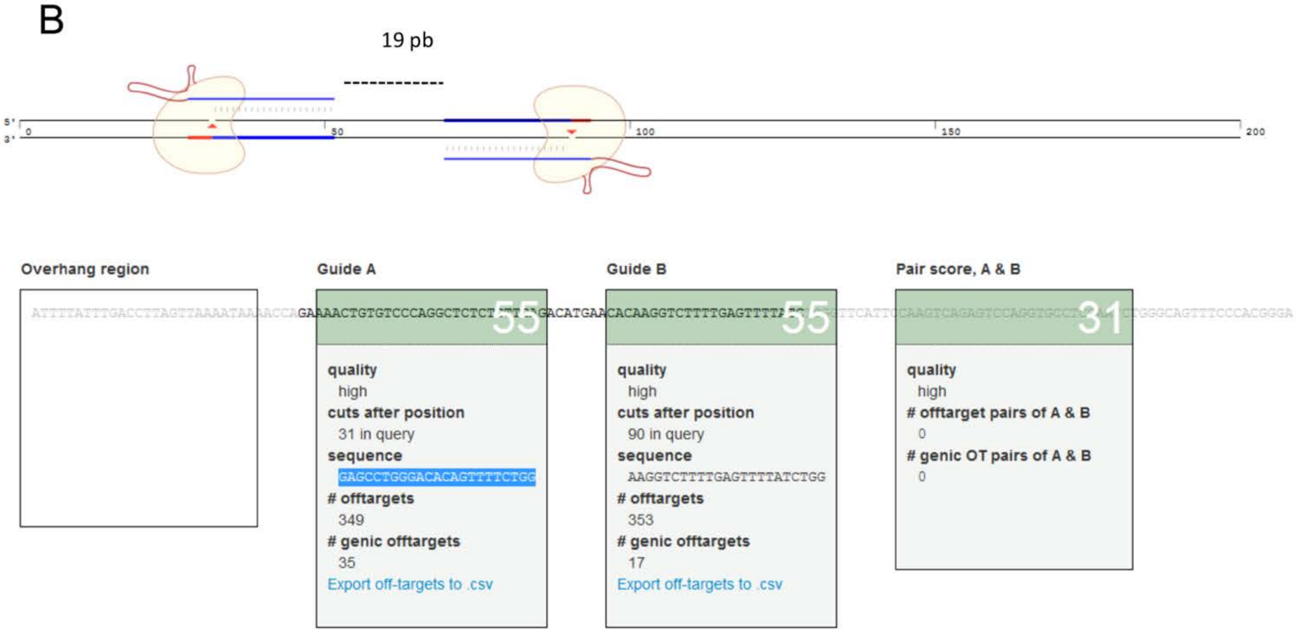


Guides 3':
gRNA NR2e3 31: GAGCCTGGGACACAGTTTTC
gRNA NR2e3 32: AAGGTCTTTTGAGTTTTATC

**Supplementary Figure 1. CRISPR/Cas9 guide position.** To perform *Nr2e3* gene edition with the Cas9 D10A nickase, we used four guides, two guides at 5' (A), and two guides at 3' (B). Selected guide sequences and position is depicted.

A

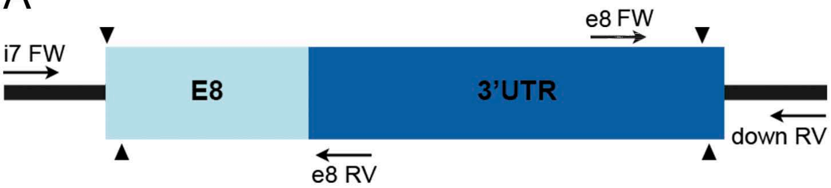


| | WT | Targeted |
|---|---|---|
| i7 FW + e8 RV | 380 bp | - |
| e8 FW + down RV | 259 bp | - |
| i7 FW + down RV | 1038 bp | 289 bp |

B 5' or upstream region

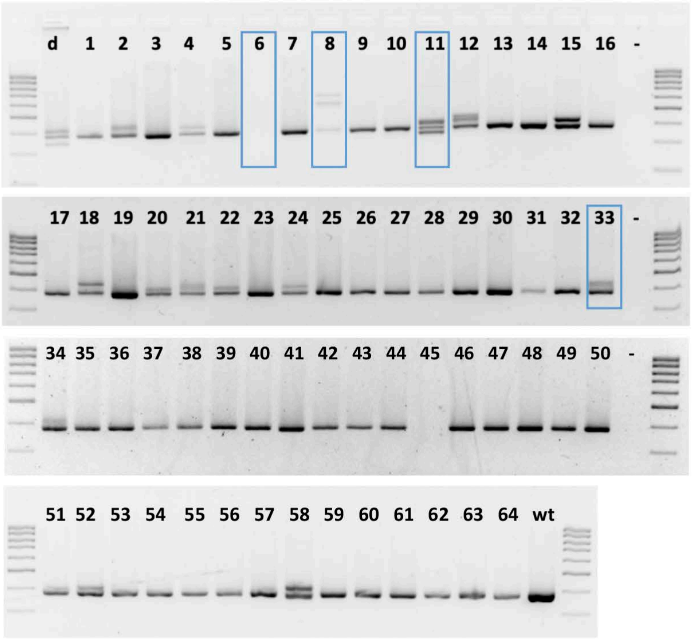


C

exon 8

**Reference WT:** TTCCTCACGGCTGAGCGCATTGAGCTTCTCTTCTTCAGAAAGACCATAGGGAACACTCCGATGGAGAAG

***Nr2e3*#02a (4/12):** TTCCTCACGGCTGAGCGCATTGAGCTTCTCTTTCTTCAGAAAGACCATAGGGAACACTCCGATGGAGAAG

***Nr2e3*#02b (3/12):** TTCCTCACGGCTGAGCGCATTGAGCTTCTCTTCTTCTTTTCTTCAGAAAGACCATAGGGAACACTCCGATGGAGAAG

***Nr2e3*#04 (3/24):** TTCCTCACGGCTG-----------------------------------------------ATGGAGAAG

***Nr2e3*#11 (7/12):** TTCCTCACGGCTGAGCGCATT---------------------------GGGAACACTCCGATGGAGAAG → Δ27

***Nr2e3*#15 (6/12):** TTCCTCAC----GAGCGCATTGAGCTTCTCTTCTTCAGAAAGACCATAGGGAACACTCCGATGGAGAAG

***Nr2e3*#16 (1/12):** TTCCTCACGGCTGAGCGCATTGAGCTTCTCTTC---AGAAAGACCATAGGGAACACTCCGATGGAGAAG

***Nr2e3*#18 (10/12):** TTCCTCACGGCTGCGGCTGAGCGCATTGAGCTTCTCTTCTTCAGAAAGACCATAGGGAACACTCCGATGGAGAAG

***Nr2e3*#20 (2/12):** TTCCTCACGGCTGAGCGCATTGAGCTTCTCTTCTTCAGAAAG------------ACTCCGATGGAGAAG

***Nr2e3*#21a (9/12):** TTCCTCACGGCTGAGCGCATTGAGCTTCTCTTCTCCAGAAAGACCATAGGGAACACTCCGATGGAGAAG

***Nr2e3*#21b (2/12):** TTCCTCACG------------------CTCTT-----------CCATAGGGAACACTCCGATGGAGAAG

***Nr2e3*#22 (3/12):** TTCCTCACG-------------AGCTTCTCTTCTTCAGAAAGACCATAGGGAACACTCCGATGGAGAAG

***Nr2e3*#24 (2/12):** TTCCTCACGGCTGAGCGCATTGAGCTTCTCTTCTTCAGAAAGACTTCAGAAAGACCATAGGGAACACTCCGATGGAGAAG

***Nr2e3*#33 (4/10):** TTCCTCACGGCTGAGCGCATTGAGCTTC**TCTTCTTCAGAAAGACC**TCTTCTTCAGAAAGACCATAGGGAACACTCCGATGGAGAAG

***Nr2e3*#34 (1/12):** TTCCTCACGGCTGAGCGCATTGAGCTTCTCTTCTTCAGAAAGACC--------------GATGGAGAAG

***Nr2e3*#48 (3/12):** TTCCTCACGGCTGAGCGCATTGAGCTTCTAGTCTTCAGAAAGACCATAGGGAACACTCCGATGGAGAAG

***Nr2e3*#52 (4/12):** TTCCTCACGGCTGAGCGCATTGAGGACCA----TTCAGAAAGACCATAGGGAACACTCCGATGGAGAAG

### D Whole deletion

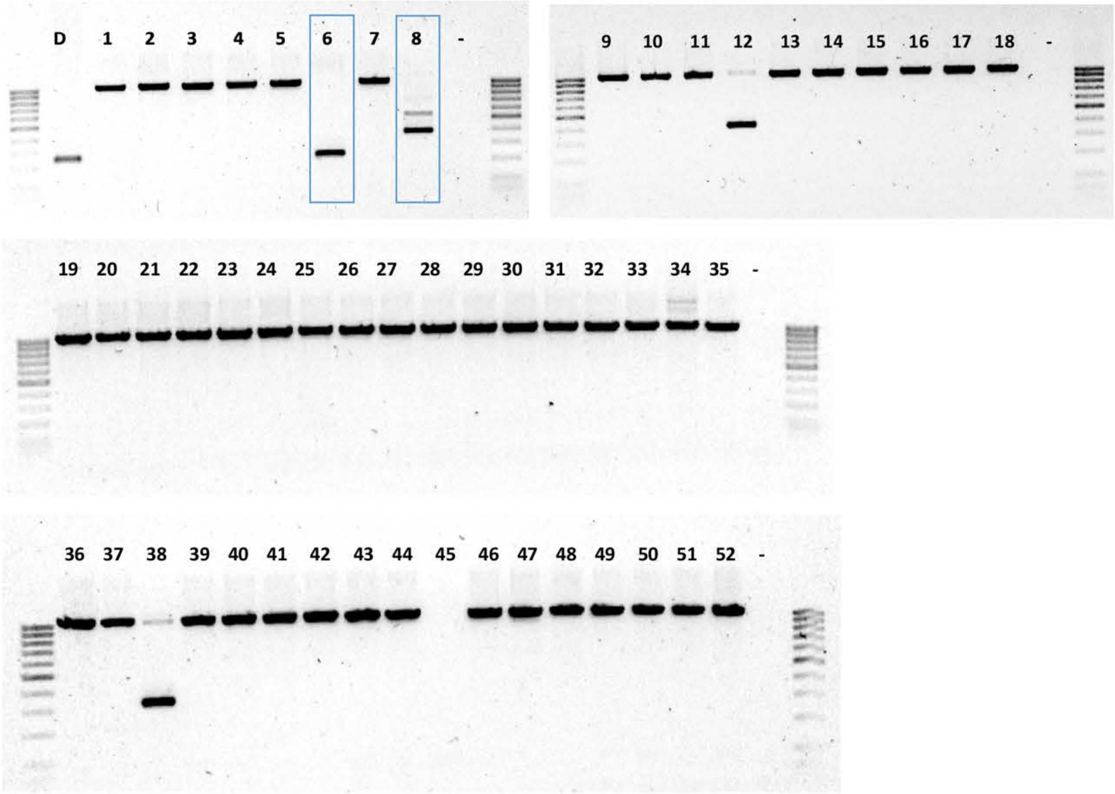


## E

**Reference WT:**

```
                                   intron 7
tggctggccctggacaactccctcccctcccccactgccttgctcggaggttggcctgacagtgtcctttctcctgttcagGTT
                                   exon 8
TGGGAAATTGCTCCTCCTGCTCCCAGTTCCTCACGGCTGAGCGCATTGAGCTTCTCTTCTTCAGAAAGACCATAGGGAACACTCC
GATGGAGAAGGTT..........................AAATAAAACCAGAAAACTGTGtcccaggctctctgttgagacatga
               downstream Nr2e3
acacaaggtcttttgagttttatctggggttcattccaagtcagag
```

***Nr2e3*#06:**

```
tggctggccctggacaactccctcccctcccctccccccactgccttgctcggaggttggcctgacagtgtc------------
------------------------------------------------------------------------------------
------------------------------------------------------------------------------------
---------------ttttgagttttatctggggttcattccaagtcagag
```

↓

ΔE8

### F 3’ or downstream region

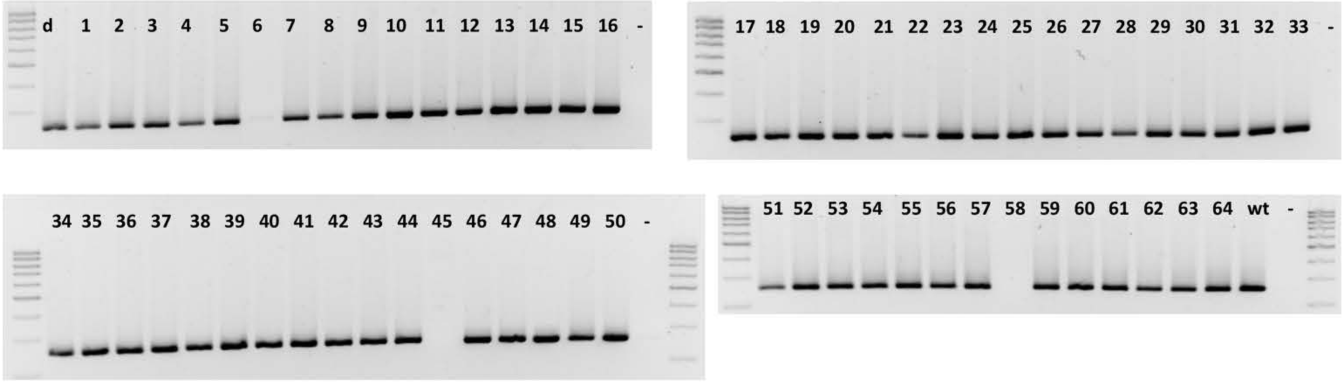

G

https://zlab.bio/guide-design-resources

| CHROM | POSITION | SEQUENCE | MISMATCH | SCORE |
|---|---|---|---|---|
| **GUIDE 51** | | | | |
| chr 9 + | 59791264 | GCTCAATGCGCTCAGCCGTGAGG | 0 | 100.00 |
| chr 15 - | 35664286 | GCTCCATGCGCTCAGCCGCGAGG | 2 | 3.84 |
| chr 13 - | 35152712 | GAAAAATGTGCTCAGCCGTGCAG | 4 | 0.84 |
| **GUIDE 52** | | | | |
| chr 9 - | 59791225 | GACCATAGGGAACACTCCGATGG | 0 | 100.00 |
| chr 15 + | 35664325 | GACCATGGGGAACAGTCCGATGG | 2 | 1.38 |
| chr 8 - | 84266017 | GGGTAAAGGGAACACTCCGAAAG | 4 | 0.79 |
| **GUIDE 31** | | | | |
| chr 9 + | 59790567 | GAGCCTGGGACACAGTTTTCTGG | 0 | 100.00 |
| chr 7 + | 36693711 | GAGTCTGGGAAACAGTTTTCAGG | 2 | 3.94 |
| chr 10 + | 102854119 | GAGTGTGTGACACAGTTTTCTAG | 3 | 2.43 |
| chr 14 - | 119837656 | GCAGCTGGGACACAGTTTTCAGG | 3 | 2.29 |
| chr 13 + | 55217157 | CAGACTGGGACTCAGTTTTCAGG | 3 | 1.42 |
| chr 8 + | 90126754 | GATCCTGGGACACTGTTTTCTGG | 2 | 1.37 |
| char2 + | 177960888 | GAGCTTGGGCGACAGTTTTCAGG | 3 | 1.3 |
| chr 11 + | 116161455 | GAGCCTCTAACACAGTTTTCAGG | 3 | 1.02 |
| **GUIDE 32** | | | | |
| chr 9 - | 59790525 | AAGGTCTTTTGAGTTTTATCTGG | 0 | 100.00 |
| chr 2 - | 162224448 | CAGGTCTTTTGAATTTTATCGAG | 2 | 3.91 |
| chr 2 + | 38667474 | GAGGTCTTAAGAGTTTTATCCAG | 3 | 1.54 |
| chr 9 + | 17578558 | AATTTCTTTTTAGTTTTATCTAG | 3 | 1.46 |
| chr 3 + | 146572367 | AAGAACTTTTAAGTTTTATCAAG | 3 | 1.45 |
| chr 12 - | 40044796 | TATCTCTCTTGAGTTTTATCCAG | 4 | 1.37 |
| chr 12 - | 40044796 | TATCTCTCTTGAGTTTTATCCAG | 4 | 1.35 |
| chr 7 + | 59967793 | AAGTTCTGTTGGGTTTTATCCAG | 3 | 1.31 |

**Supplementary Figure 2. Generation of *Nr2e3* mutant alleles by gene editing with CRISPR-Cas9D10A nickase. A)** Schematic representation of *Nr2e3* exon 8 targeting. Arrowheads indicate the position of the targeting sequences for the four CRISPR guide RNAs. The position of the genotyping PCRs is shown by black arrows. The predicted size of the PCR products for the wild type and targeted mutant alleles is indicated within the box. **B)** Genotyping of gene-edited alleles at the 5' position (junction of intron 7 and exon 8) showing the PCR products. The amplification of additional PCR bands indicated mosaic mice carrying gene-edited alleles. Blue boxes indicate mice selected for further analysis. Lane 11 corresponds to the mosaic mouse carrying the Δ27 mutant allele. **C)** Sequence of PCR bands amplified from gene-edited animals showing different *Nr2e3* gene-edited alleles at the 5'-end. Many modified alleles with partial deletions and modifications at the junction of intron7- exon 8 were generated. In red, PAM sequences; in blue, additional nucleotides added after DNA repair of the DSB. Dashes indicate deleted sequences, whereas the sequence in bold indicates a duplicated sequence in tandem. Yellow squares indicate Δ27 allele. **D)** Genotyping of gene-edited alleles for full exon 8 deletion, showing the PCR products. Blue boxes indicate interesting modified alleles. Lane 6 corresponds with the ΔE8 mutant. **E)** Sequence of the PCR band amplified from gene-edited animals with the complete targeted deletion. Only 1 of the 64 microinjected mice carried the complete exon 8 deletion. In red, PAM sequences; in blue, additional nucleotides added after DNA repair of the DSB. Dashes indicate deleted sequences, whereas the sequence in bold indicates a duplicated sequence in tandem. Yellow squares indicate ΔE8 allele. **F)** Genotyping of gene-edited alleles at the 3' position showing the PCR products. **G)** List of on-target (100% match) and potential off-target regions (with very low scores) for each guide, which were tested in all the pups. No off-target events were detected.

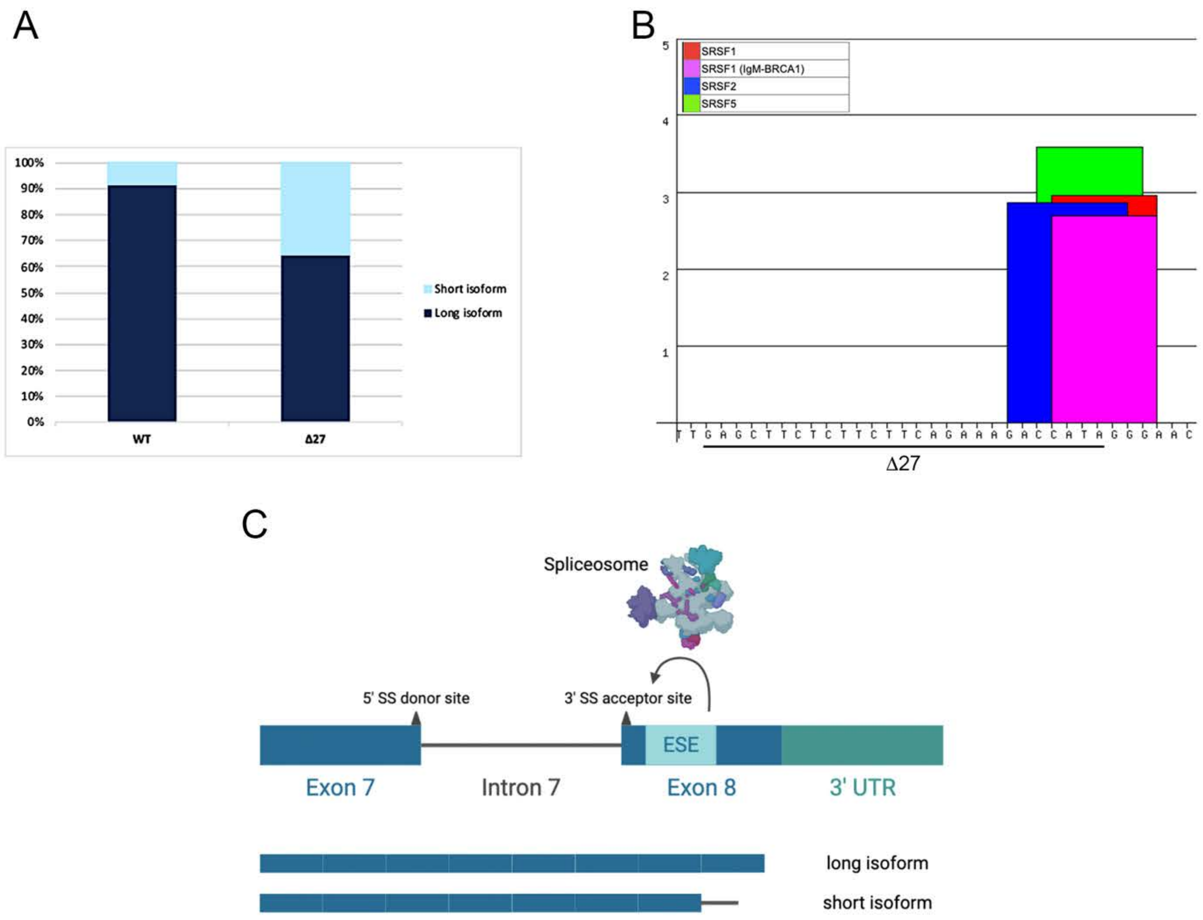


**Supplementary Figure 3. Identification of a putative ESE located in *Nr2e3* exon 8**.
**A)** Relative quantification of *Nr2e3* isoform expression indicating the percentile of each isoform with respect to total *Nr2e3* expression set to 100%. In adult retinas, wild type allele produce ~92% of long isoform and ~8% of short isoform, whereas Δ27 allele produces ~64% of long isoform and ~36% of short isoform. **B)** Graphic of the *in silico* prediction by ESE Finder (http://rulai.cshl.edu) that identifies a putative ESE located within the 27 nucleotides deleted in the Δ27 mutant. Bars indicate predicted binding sites for specific SR proteins (SRSF1 in red and pink, SRSF2 in blue and SRSF5 in green). The deletion of these 27 nt completely ablates this ESE. **C)** Schematic representation of the enhanced splicing of intron 7 by the ESE in exon 8, according to the *in silico* predictions and our *in vivo* results. SR proteins recognizing and binding the ESE in exon 8 would recruit splicing factors that stimulate splicing of intron 7, thus favoring the production of the long isoform. The short isoform is generated when intron 7 is retained, which is more frequent in the Δ27 allele due to the deletion of the ESE motif.

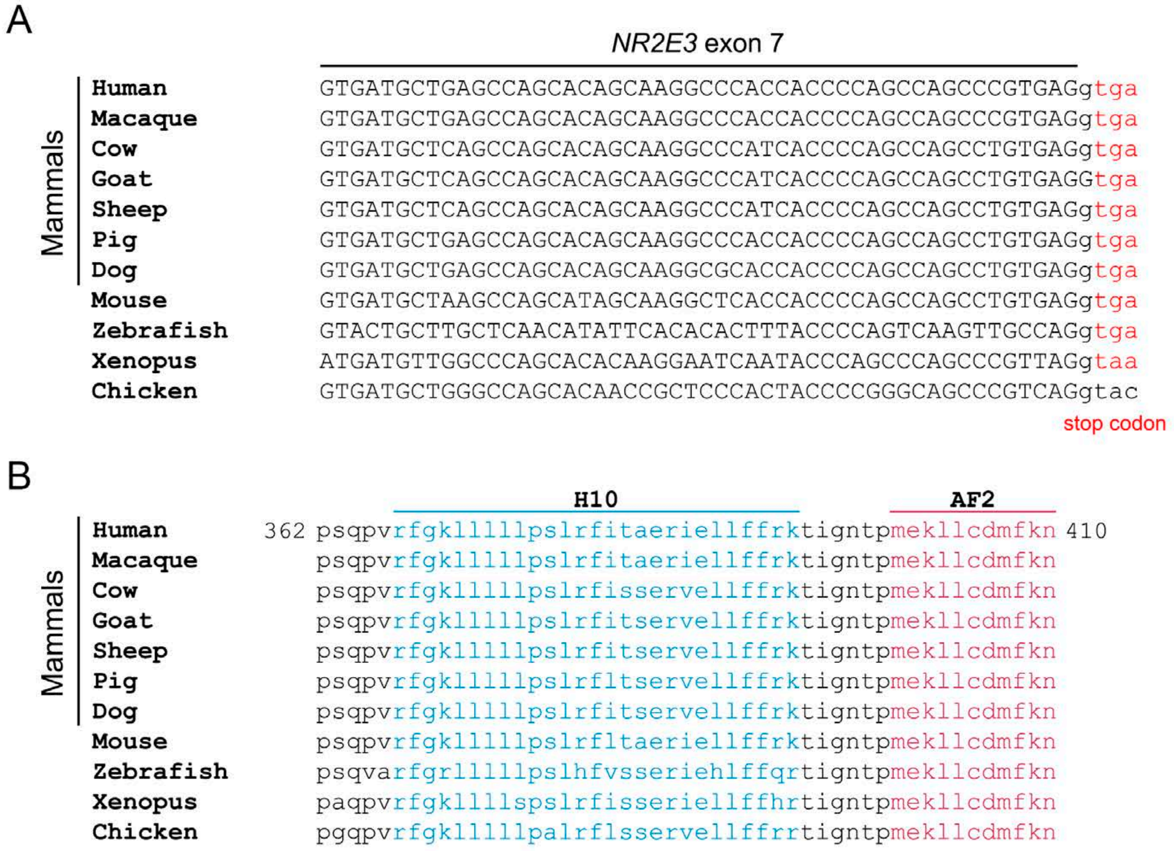


**Supplementary Figure 4. Phylogenetic alignment of *Nr2e3* DNA and protein sequences in vertebrate species. A)** Comparative alignment of the *Nr2e3* DNA sequence at the junction between the end of human exon 7 (upper case letters) and the beginning intron 7 (lower case letters). The frame of exon 7 ends with the two first nucleotides of an Arginine codon. In the transcript isoforms where intron 7 is retained, the NR2E3 protein would end in the STOP codon encoded in the following triplet (nucleotides in red), which is evolutionarily conserved among species. **B)** Comparative alignment of the NR2E3 protein sequence, which is extremely conserved among species. H10 and AF2 domains are indicated in light and dark blue, respectively. The first amino acid in H10 (Arginine) is encoded by the last two nucleotides of exon 7 and the first nucleotide in exon 8. In the shorter protein isoform, this Arginine residue is the C-terminal amino acid.

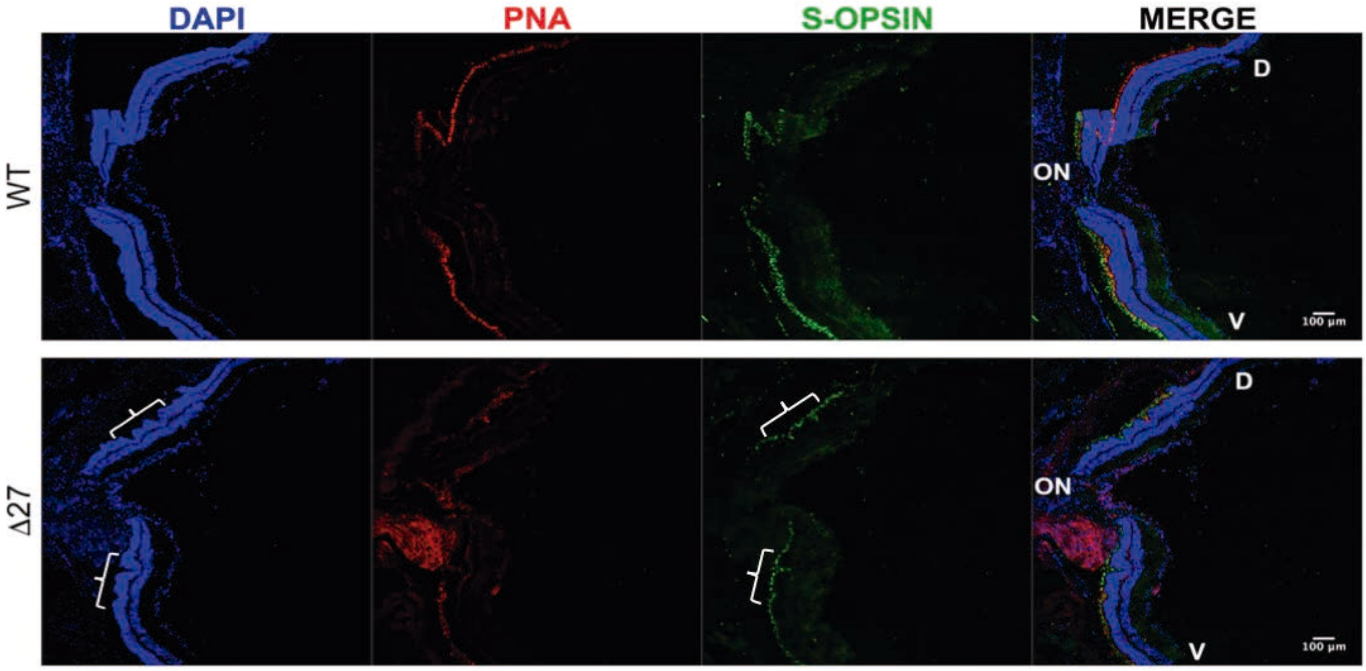


**Supplementary Figure 5. Δ27 mutant retinas show distinct S-cone distribution.** S-cones are uniformly distributed in the retina of the Δ27 mutants. Retinal sections centered at the optic nerve (ON) were stained for detection of nuclei (DAPI, blue), all type of cones (PNA, red) and S-opsin (S-cones, green). S-cones appear to be uniformly distributed in the Δ27 homozygous compared to the wildtype retina, where it follows a dorsal-ventral gradient (D-V), being the highest concentration in the ventral (V) retina. Note the cone-rich invaginations in the central retina of the Δ27, indicated by white brackets. Scale bar: 100 µm.
